# Cardiomyocyte-specific deletion of PTP1B protects against HFD-induced cardiomyopathy through direct regulation of cardiac metabolic signaling

**DOI:** 10.1101/2023.05.19.541546

**Authors:** Yan Sun, Vasanth Chanrasekhar, Chase W. Kessinger, Peiyang Tang, Yunan Gao, Sarah Kamli-Salino, Katherine Nelson, Mirela Delibegovic, E. Dale Abel, Maria I. Kontaridis

**Author notes:** Corresponding author: Maria I. Kontaridis, Ph.D., Gordon K. Moe Professor and Chair, Biomedical Research and Translational Medicine Masonic Medical Research Institute, Associate Professor of Medicine, Part-time Beth Israel Deaconess Medical Center Harvard Medical School, 2150 Bleecker St, Utica, NY 13501.

## Abstract

**Background:** Heart failure is the number one cause of death worldwide and mortality is directly correlated with the high incidence of obesity and diabetes. Indeed, the epidemic phenomenon of obesity was projected to reach 50% in the US by the year 2030. However, the mechanisms linking metabolic dysfunction with heart disease are not clear. Protein Tyrosine Phosphatase 1B (PTP1B), a negative regulator of insulin signaling, is considered to be an emerging therapeutic target against the development of obesity, insulin resistance, and diabetes. Increased PTP1B levels and activity have been observed in brain, muscle and adipose tissues isolated from obese and/or diabetic animals, as well as in human obese human patients. Its role, however, and the mechanisms by which it modulates metabolic processes in the heart remain unknown.

**Method and Results:** We generated cardiomyocyte (CM)-specific PTP1B knock-out (PTP1B^fl/fl^::ꭤMHC^Cre/+^) mice to investigate the cardiomyocyte-specific role of PTP1B in response to high fat diet (HFD)-induced cardiac dysfunction. While we did not observe any physiological or functional cardiac differences at baseline, in response to HFD, we found that PTP1B^fl/fl^::ꭤMHC^Cre/+^ mice were protected against development of cardiac hypertrophy, mitochondrial dysfunction, and diminished cardiac steatosis. Metabolomics data revealed that hearts with CM-specific deletion of PTP1B had increased fatty acid oxidation and NAD^+^ metabolism, but reduced glucose metabolism; we further validated these findings by real-time qPCR analysis. Mechanistically, we identified a novel PTP1B PKM2-AMPK axis in the heart, which acts as a molecular switch to promote fatty acid oxidation. In this regard, we identified that hearts from PTP1B^fl/fl^::ꭤMHC^Cre/+^ mice had upregulated levels of nicotinamide adenine dinucleotide (NAD^+^) and NAD phosphate (NADPH), leading to higher levels of nicotinamide phosphoribosyl transferase (NAMPT), the rate-limiting step of the NAD^+^ salvage pathway and an enzyme associated with obesity and diabetes.

**Conclusions:** Together, these results suggest that CM-specific deletion of PTP1B mediates a substrate switch from glucose to fatty acid metabolism, protecting hearts against development of HFD-induced cardiac hypertrophy and dysfunction through mechanisms involving a novel PTP1B/PKM2/AMPK axis that is critical for the regulation of NAMPT and NAD^+^ biosynthesis.

## Introduction

Obesity has reached epidemic proportions worldwide and continues to rise at an alarming rate. Projections from the World Health Organization suggest that 50% of the US will be classically defined as obese by the year 2030, with the most jeopardized demographic being in children^1^. Worse, obesity increases the risk for developing multiple diseases, including type 2 diabetes, certain cancers, and cardiovascular disease (CVD)^2^. With regards to the latter, obesity mediates adverse effects on glucose and lipid levels, increases arterial blood pressure, induces inflammation, and reduces pulmonary function, characteristics that can lead to cardiac hypertrophy, dysfunction and/or heart failure if left untreated ^3, 4^ ^5, 6^. Unfortunately, despite knowing the contributing factors that lead to obesity-related CVD, the molecular mechanisms for how obesity and the consequent associated physiological changes directly lead to CVD remain poorly understood.

Cardiac metabolism is complex, involving hormonal regulation and activation of multiple downstream signaling pathways that control the balance between glucose and fatty acid utilization^7^. Under normal physiological conditions, about 95% of the adenosine triphosphate (ATP) generated and utilized by the heart comes from the process of oxidative phosphorylation (OXPHOS) in the mitochondria. Mitochondrial OXPHOS is fueled by the energy garnered from fatty acid oxidation (FAO), the tricarboxylic acid (TCA) cycle, and to a lesser extent, pyruvate dehydrogenase; the remaining 5% of ATP is supplied by the oxidation of glucose, lactate and ketone bodies^8, 9^.

In contrast, cardiac substrate metabolism switches to favor glucose metabolism over FAO in response to hemodynamic stress and pathological stimuli^8^. Specifically, in response to obesity and high-fat diet (HFD), the heart undergoes marked alterations in energy metabolism, increasing fatty acid uptake, enhancing oxidation as a result of increased supply of fatty acids ^10–12^, and contributing to lipotoxic heart disease, which may increase the risk of heart failure. However, the mechanisms governing excess lipid accumulation and adverse sequelae in cardiomyocytes in response to HFD entails more comprehensive investigation.

Systemic metabolic substrate preference is largely under the control of the circulating hormone insulin. Indeed, insulin signaling is a critical modulator of the body’s ability to metabolize carbohydrates, lipids and proteins^13^, and is integral in regulating homeostatic processes that control cellular proliferation^14^, differentiation^15^, and apoptosis^16^ . Briefly, in the presence of insulin, the insulin receptor (IR) phosphorylates insulin receptor substrate (IRS) proteins, inducing the activation of critical downstream pathways that include the phosphatidylinositol 3-kinase (PI3K)–protein kinase B/AKT (AKT) pathway, which plays a major role in metabolic actions of insulin, and the Ras–mitogen-activated protein kinase (MAPK) pathway, which regulates gene transcription and cooperates with the PI3K pathway to regulate cell growth and differentiation^17–19^.

The protein tyrosine phosphatase non-receptor type 1 (*PTPN1*), which encodes the non transmembrane protein tyrosine phosphatase (PTP1B), is a ubiquitously expressed protein that functions as a negative regulator of insulin signaling^20–22, 23^. Because of this, PTP1B is considered an emerging potential therapeutic target against the development of obesity, insulin resistance, and diabetes. Indeed, increased PTP1B levels and activity have been observed in brain, muscle and adipose tissues isolated from obese and/or diabetic animals^24–28^, as well as in human obese patients^24, 29–31^. Conversely, mice with germline deletion of PTP1B are resistant to obesity and have increased insulin sensitivity in response to HFD^32^. Interestingly, neither liver-specific nor adipose-specific deletion of PTP1B affect weight gain in HFD-fed mice ^21, 33^, suggesting a non-autonomous regulation of metabolism and/or obesity by PTP1B. In this regard, mice with neuronal-specific or pro-opiomelanocortin (POMC)-specific PTP1B deletion exhibit decreased body weight in response to HFD, effects found to be associated with increased leptin sensitivity and improved glucose homeostasis^34, 35^.

Recent studies have shown that in addition to insulin signaling, PTP1B dephosphorylates and activates pyruvate kinase muscle isozyme 2 (PKM2) in pancreatic cancer cells and cultured adipocytes, inducing glycolysis by promoting the conversion of phosphoenolpyruvate to pyruvate^33, 36^. Activated PKM2 also affects the adenosine monophosphate (AMP)/ATP ratio by inhibiting AMP-activated protein kinase (AMPK) activity^36, 37^, thereby potentially controlling metabolic and cellular energy homeostasis by blocking FAO^38, 39^ and increasing lipogenesis^40^. Additionally, AMPK controls nicotinamide phosphoribosyl-transferase (NAMPT) expression^41^, the rate-limiting enzyme in the nicotinamide adenine dinucleotide (NAD) salvage biosynthesis pathway that is responsible for converting nicotinamide (NAM) to nicotinamide ribonucleotide (NMN)^42^. This PKM2/AMPK axis may be regulated by PTP1B, but how and whether this is modulated specifically in the heart remains unknown.

Given the complexities of PTP1B in both insulin/AKT and PKM2/AMPK signaling, it is critical to determine the direct role of PTP1B in the heart. It has been previously established that increased PTP1B activity is associated with increased incidence of heart failure in both rats and humans^43^. Genome wide expression analysis studies demonstrate that increased pressure overload in the heart increases expression of PTP1B^43^. Both cardiac contractile and intracellular calcium cation dysfunction are also associated with elevated expression levels of PTP1B^44^. Conversely, endothelial-specific deletion of PTP1B protects against pressure overload-induced heart failure; inhibition of PTP1B in these cells drives activation of vascular endothelial growth factor (VEGF) signaling and angiogenesis, inducing migration and proliferation of microvascular endothelial cells, reducing hypoxia, and preventing fibrosis ^45, 46^.

Taken together, we believe PTP1B is an integral signaling protein involved in metabolism and a nodal enzyme critical for the regulation of cardiac function. However, a direct role for PTP1B in cardiac insulin resistance in cardiomyocytes (CMs) and its effect on obesity induced pathological cardiac remodeling remains as yet unknown. Here, we investigated the role PTP1B directly in CMs in response to HFD to assess whether deletion of this enzyme in the heart may protect against the development of obesity-induced cardiomyopathy.

## Materials and methods

### Mice

To generate mice with cardiomyocyte-specific PTP1B knock-out (PTP1B^fl/fl^::ꭤMHC ^Cre/+^), C57Bl6 mice with loxP-flanked PTP1B floxed alleles (PTP1B^fl/fl^) (courtesy of Benjamin G. Neel, NYU Grossman School of Medicine, Laura and Isaac Perlmutter Cancer Center, New York; ^1^) were mated with transgenic α myosin heavy chain (*MYH6*) mice that drive the expression of Cre (αMHC^Cre/+^) (generously provided by Dr. Dale Abel, University of California, Department of Medicine, Los Angeles; ^47^). Mice were weaned and placed on either normal diet (ND) (PicoLab Rodent Diet20#5053) or HFD (Research Diet #12492, 60 kcal% fat) at post-natal day 30, for a period of 10 weeks (males) or 20 weeks (females). NOTE: Unlike the males, which showed a significant cardiac phenotype by 10 weeks on HFD, the female mice did not have an effect in this time period, irrespective of genotype; therefore, our female animals were studied over 20 weeks, an extended time course, to reveal the phenotype in these mice. Otherwise, we used age-matched mice for all experimental groups. All mice were on the C57Bl6 background, and the αMHC^Cre/+^ mice were used as our control group. All animal procedures were approved by the Masonic Medical Research Institute Animal Care and Use Committee. Our PHS assurance number is D16-00144(A3228-01) and we are an AAALAC (#001865) accredited institution.

### RNA Extraction and Real-time PCR

RNA was isolated using Trizol (Invitrogen) and purified with an RNeasy kit (Qiagen). A total of 1 μg RNA was reverse-transcribed with Iscript Supermix (Bio-rad). cDNA was then used to amplify the target genes by SYBR Green PCR Master Mix (Thermo Fisher Scientific). 18S ribosomal RNA *(18S),* eukaryotic translation elongation factor 1 (*EEF1A1*) and ribosomal protein L4 (*RPL4*) were used as our control housekeeping genes. Data were quantified using the comparative C^T^ method (ΔΔCT). For primer sequences and PCR conditions, please see the supplemental information (Table S1).

### Biochemical Studies

Tissue lysates were prepared by homogenizing the tissue in radioimmunoprecipitation (RIPA) buffer (25 mmol/l Tris-HCl [pH 7.4], 150mmol/l NaCl, 0.1% SDS, 1% NP-40, 0.5% sodium deoxycholate, 5mmol/l EDTA), 1mmol/l sodium fluoride, 1mmol/l sodium orthovanadate, and a protease cocktail at 4°C, followed by sonication. For immunoblots, proteins were resolved by sodium dodecyl-sulfate polyacrylamide gel electrophoresis (SDS-PAGE) and transferred to nitrocellulose membranes. Immunoblots were performed with anti-PTP1B (Abcam), anti-AKT, anti-phospho-AKT (Ser473), anti-extracellular signal-regulated kinase (ERK) 1/2, anti-phospho-ERK 1/2 (Thr202/Tyr204), anti-AMPK, anti-phospho-AMPK (Ser485), anti microtubule-associated protein 1A/1B light chain 3B (LC3B), anti-PKM2, anti-phospho-PKM2 (Tyr105) (Cell Signaling Technology) or anti glyceraldehyde 3-phosphate dehydrogenase (GAPDH) (Santa Cruz Biotechnology, Inc).

### Histology

Hearts were isolated and fixed in 4% paraformaldehyde for 24 hours and then paraffin embedded. Sections were stained with hematoxylin/eosin (H&E) or Masson’s trichrome and images were taken using a Keyence BXZ microscope. Cryosections (8 µm) of hearts were permeabilized with 0.1% Triton X in PBS for 5 mins at room temperature, hearts were blocked with normal horse serum for 30 mins. Coverslips were incubated with wheat germ agglutinin (WGA)-Alexa Fluor 647 (1:100) (Thermo Fisher Scientific) antibodies in a humid chamber. Slides were imaged by a Zeiss Confocal microscope. The quantitative analysis of CM area size included 5 different fields from four mice per group and 3 sections per heart. The total number of myocytes counted was 1-2x10^3^ cells per mouse.

### Echocardiography

Transthoracic echocardiography was conducted on non-anesthetized animals as described previously^2^, using a Visual Sonics Vevo 3100® high-frequency ultrasound rodent imaging system. Hearts were imaged in the two-dimensional parasternal short axis view, and an M-mode echocardiogram of the mid ventricle was recorded at the level of papillary muscles. Heart rate, posterior wall thickness (LVPW), and end-diastolic and end-systolic internal dimensions of the left ventricle (LVIDd and LVIDs, respectively) were measured from the M-mode image.

### Mitochondrial Respiration

Mitochondria were isolated from the heart as previously described^48^. Briefly, hearts were dissected and washed in ice-cold mitochondrial isolation buffer. Tissues were cut into small pieces and homogenized with Potter-Elvehjem tissue grinder. Tissue pieces were disrupted with at least 30 strokes, and then centrifuged at gradient speed to obtain a pellet. The pellet was resuspended in mitochondrial assay solution containing 10 mM glutamate and 5 mM malate. Mitochondrial respiration was determined using an XF96 Extracellular Flux Analyzer (Seahorse Bioscience). For those measurements, 50 µg mitochondria were plated and centrifuged at 2,000 g for 20 mins to promote adherence. Oligomycin (1µg/mL) and carbonyl cyanide-p-trifluoromethoxyphenyl-hydrazon (FCCP) (20mM) were used to inhibit ATP synthase by blocking its proton channel. Rotenone and Antimycin (40mM) were used to inhibit Complex I and Complex III-dependent respiration. All readings were normalized to protein content. N=6 mice per group.

### Detection of Mitochondrial Membrane Potential

JC-1 (Abcam, USA) was used to assess the mitochondrial membrane potential. Isolated cardiomyocytes were incubated with 5µM JC-1 staining solution at 37 ℃ free light for 10 min according to manufacturer’s instruction. The adult cardiomyocytes were washed by culture medium and imaged by a Zeiss Confocal microscope. JC1 monomers and aggregates were both excited at 488 nm. Detection of fluorescence for JC1 monomers and aggregates were performed respectively at 530 and 590 nm. Ratio F(aggregate)/F(monomer) was subsequently evaluated using Image J^3^.

### Transmission Electron Microscopy (TEM)

TEM studies were performed by the SUNY Upstate University Transmission Electron Microscopy Center. Mice were sacrificed and their hearts were perfused with 2.5% (vol/vol) glutaraldehyde in 0.1M Na-cacodylate buffer (pH 7.4). The hearts were immediately cut into pieces < 1mm^3^ and fixed in the 2.5% (vol/vol) glutaraldehyde in 0.1M Na-cacodylate buffer (pH 7.4) for 2 hours. Following fixation with 1% osmium tetroxide in 0.1M cacodylate buffer, the tissue was dehydrated in a graded series of ethanol and embedded in Spurr’s resin. Semithin (0.5μm) and ultrathin (90nm) sections were cut, mounted on copper grids, and post-stained with uranyl acetate and lead citrate. Sections were viewed with a JEM 2100F transmission electron microscope.

### Adult Primary Cardiomyocyte Isolation

Mice were injected with intraperitoneal heparin (40 unit/mice), and their hearts were isolated and perfused via the aorta. The perfusion buffer consisted of KCl (14.7mM), NaCl (120.4mM), KH_2_PO_4_ (0.6mM) Na_2_HPO_4_ (0.6mM), MgSO_4_7H_2_O (1.2mM), 2,3 butanedione monoxime (10mM), taurine (30mM), HEPES (10mM), and glucose (5.5mM). The heart was digested with collagenase II digestion buffer (2 mg/ml) for approximately 8-10 min. The heart was cut from the cannula and placed in the dish with digestion buffer and stopping buffer [12.5µM CaCl_2_ and 10% exosome depleted FBS in perfusion buffer]. Isolated cardiomyocytes were then cultured in a six well plate (treated with laminin), with myocyte culture medium [ScienCell #6201], 5% exosome depleted FBS, and 1% penicillin/streptomycin.

### Metabolomics Studies (as reported by Metabolon, Inc)

#### Sample preparation

Metabolomic analysis was performed by Metabolon, Inc. Briefly, heart samples were immediately snap-frozen in liquid nitrogen and shipped overnight on dry ice to the Metabolon, Inc. Samples were prepared using the automated MicroLab STAR® system from Hamilton Company. Several recovery standards were added prior to the first step in the extraction process for QC purposes. To isolate the protein, small molecules bound to protein or trapped in the precipitated protein matrix were dissociated. To recover chemically diverse metabolites on these complexes, proteins were precipitated with methanol under vigorous shaking for 2 min (Glen Mills GenoGrinder 2000), followed by centrifugation. The resulting extract was divided into five fractions: two for analysis by two separate reverse phases/UPLC-MS/MS methods with positive ion mode electrospray ionization (ESI), one sample for analysis by RP/UPLC-MS/MS with negative ion mode ESI, one fraction for analysis by HILIC/UPLC-MS/MS with negative ion mode ESI, and one sample was reserved for backup. Samples were placed briefly on a TurboVap® (Zymark) to remove the organic solvent. The sample extracts were stored overnight under nitrogen before preparation for analysis.

#### QA/QC

Several types of controls were analyzed in concert with the experimental samples: a pooled matrix sample generated by taking a small volume of each experimental sample served as a technical replicate throughout the data set; extracted water samples served as process blanks; and a cocktail of QC standards, carefully chosen not to interfere with the measurement of endogenous compounds, were spiked into every analyzed sample. These controls allowed for optimal instrument performance monitoring and aided the chromatographic alignment. Instrument variability was determined by calculating the median relative standard deviation (RSD) for the standards that were added to each sample prior to injection into the mass spectrometer. Overall process variability was determined by calculating the median RSD for all endogenous metabolites (i.e., non-instrument standards) present in 100% of the pooled matrix samples. Experimental samples were randomized across the platform run with QC samples spaced evenly among the injections.

#### Metabolite Quantification and Data Normalization

Peaks were quantified using the area under-the-curve. For studies spanning multiple days, a data normalization step was performed to correct variation resulting from instrument inter-day tuning differences. Essentially, each compound was corrected in run-day blocks by registering the medians to equal one (1.00) and normalizing each data point proportionately (termed the “block correction”). For studies that did not require more than one day of analysis, no normalization was necessary, other than for purposes of data visualization. In certain instances, biochemical data may have been normalized to an additional factor (e.g., cell counts, total protein as determined by Bradford assay, osmolality, etc.) to account for differences in metabolite levels due to differences in the amount of material present in each sample.

### NAD^+^ and NADH Measurements

NAD^+^ and NADH were measured using the EnzyChrom^TM^ NAD^+^/NADH Assay Kit according to the manufacturer’s protocol (ECND-100, Bioassay Systems, Hayward, CA). Briefly, 20mg of heart tissue was weighed out for each sample and washed with cold PBS. Samples were homogenized in a 1.5mL Eppendorf tube with either 100μL NAD extraction buffer for NAD determination or 100μL NADH extraction buffer for NAD determination. Extracts were heated at 60°C for 5 min and neutralized by addition of 20µL Assay Buffer and 100µL of Neutralizing Buffer. Samples were vortexed and spun down at 14,000 rpm for 5 min. Supernatants were then collected and used for the NAD/NADH assay. Here, 40μL of the supernatant was added to 80μL of the Working Reagent and optical density at 565nm was read at time zero and after a 15-min incubation at room temperature.

### Statistical Analysis

All values in graphs are expressed as the mean ± SEM. Normality was tested with the D’Agostino-Pearson normality test. If the data showed a normal distribution, pairwise testing was performed with the student’s *t* test or multiple group comparisons were performed by 2-way ANOVA, followed by post-hoc Bonferroni correction (GraphPad Prism 9).

## Results

### Generation of cardiomyocyte-specific PTP1B knock-out mice

To investigate the role of PTP1B in obesity-induced cardiomyopathy, we generated a mouse model with CM-specific deletion of PTP1B. We crossed PTP1B^fl/fl^ mice^34^ with mice expressing Cre recombinase under the control of the α-MHC promoter to generate PTP1B^fl/fl^::ꭤMHC^Cre/+^ mice, as well as their background controls, PTP1B^+/+^::ꭤMHC^Cre/+^ mice (Figure S1A, B). While PTP1B protein expression was detected normally in male and female PTP1B^+/+^::ꭤMHC^Cre/+^ mice, PTP1B was undetectable in CMs isolated from PTP1B^fl/fl^: ꭤMHC^Cre/+^ mice (Figure S1C). PTP1B expression in other cardiac cell types and/or in other tissues, including the spleen, remained unchanged (Figure S1D, S1E). Male and female PTP1B^+/+^::ꭤMHC^Cre/+^ and PTP1B^fl/fl^:: ꭤMHC^Cre/+^ mice were born at expected mendelian ratios and did not display any abnormal phenotypic features (Table S2 and data not shown).

### Cardiomyocyte-specific deletion of PTP1B prevents HFD-induced cardiomyopathy

To investigate the role of obesity-associated PTP1B in CMs, we examined the consequence of PTP1B^+/+^::ꭤMHC^Cre/+^ and PTP1B^fl/fl^:: ꭤMHC^Cre/+^ mice in response to a HFD diet regime. Under conditions of ND, hearts of PTP1B^+/+^::ꭤMHC^Cre/+^ and PTP1B^fl/fl^:: ꭤMHC^Cre/+^ mice were of similar size and weight for both males and females (Figure 1A, 1B, S2A, S2B), suggesting that PTP1B has minimal physiological effects on the heart at baseline. Following a regiment of HFD for 10 weeks in male mice, however, we found that control hearts developed hypertrophy with a 1.2 fold change in heart weight to tibia length, whereas hearts from PTP1B^fl/fl^:: ꭤMHC^Cre/+^ mice remained normal, suggesting that deletion of PTP1B in CMs protected against HFD-induced cardiac hypertrophy in males (Figure 1A, 1B). Interestingly, HFD in females did not significantly impact overall heart size in either PTP1B^+/+^::ꭤMHC^Cre/+^ or PTP1B^fl/fl^:: ꭤMHC^Cre/+^ mice after 10 weeks HFD (data not shown). To determine if phenotypic differences could be demonstrated at later time points in the females, we continued HFD treatment in this group for an additional 8 weeks, to 20 weeks total (Figure S2A, S2B). Even at this later time point, we did not observe significant phenotypic differences in the females in response to HFD in either genotype, suggesting that female mice are generally more protected against HFD induced cardiac hypertrophy, as has been suggested by previous studies ^49, 50^.

**Figure 1.**
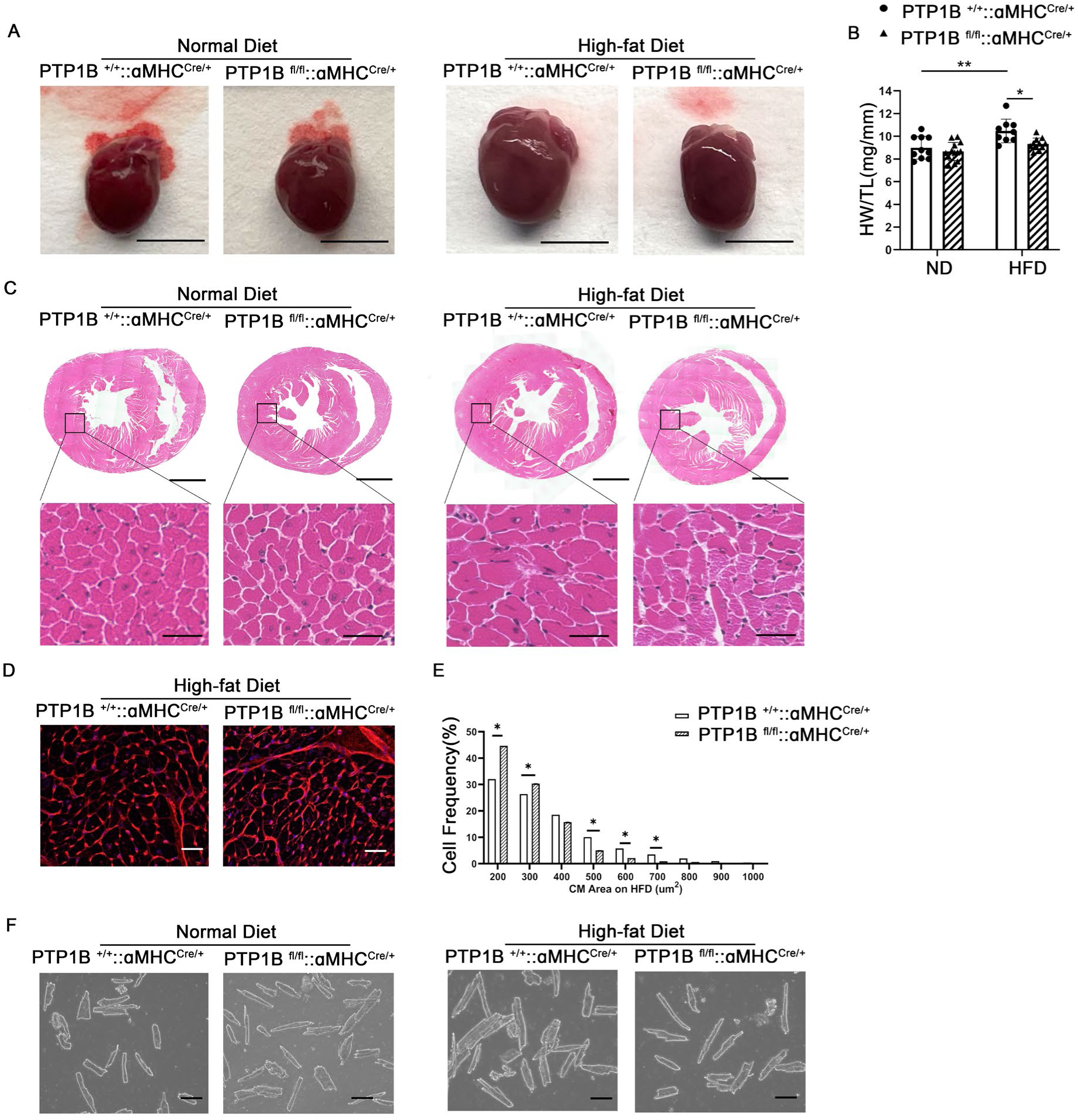
Cardiomyocyte-specific deletion of PTP1B prevents HFD-induced cardiomyopathy. **A.** Representative photographs of hearts from male PTP1B^+/+^::ꭤMHC ^Cre/+^ and PTP1B^fl/fl^::ꭤMHC ^Cre/+^ mice maintained on a normal diet (ND) or high-fat diet (HFD) for 10 weeks, scale bar=50 mm. **B**. Heart weight to tibia length ratios from male control or PTP1B^fl/fl^::ꭤMHC ^Cre/+^ mice on ND or HFD. n= 8-12/group. * p<0.05 and **p<.01 from two way ANOVA. **C**. Representative H&E whole heart cross-sections from PTP1B^+/+^::ꭤMHC ^Cre/+^ and PTP1B^fl/fl^::ꭤMHC ^Cre/+^ male mice maintained on ND or HFD for 10 weeks. Upper panel scale bar=1mm; lower panel scale bar=50µm. **D**. Wheat germ agglutinin (WGA) staining (red) of heart cross-sections from control and PTP1B^fl/fl^::ꭤMHC ^Cre/+^ male mice fed a ND or HFD for 10 weeks. Scale bar= 50µm. **E**. Frequency distribution of cardiomyocyte (CM) area from control or PTP1B^fl/fl^::ꭤMHC ^Cre/+^male mice fed ND or HFD. N=4 mice per group, with at least 1x10^3^ cell counts per heart. Data are represented as mean ± *S*EM, * p<0.05 based on Student’s *t*-test. **F**. Representative photomicrograph of ventricular myocytes isolated from PTP1B^+/+^::ꭤMHC ^Cre/+^ and PTP1B^fl/fl^::ꭤMHC ^Cre/+^ male mice maintained on ND or on HFD. Scale bar= 50 µm.

Despite this result, when analyzing the heart histologically by H&E staining, we found that HFD increased overall CM cell size, induced myocardial disarray and showed enlarged nuclei in both male and female PTP1B^+/+^::ꭤMHC^Cre/+^ mice (Figure 1C, S2C). Deletion of PTP1B in hearts isolated from both male and female mice showed near normal myocardial size and structure (Figure 1C, S2C), suggesting that deletion of PTP1B protects hearts from HFD-induced hypertrophy. Similarly, HFD-treated PTP1B^+/+^::ꭤMHC^Cre/+^ hearts displayed increased CM cross-sectional area of left ventricular CMs as compared to PTP1B^fl/fl^:: ꭤMHC^Cre/+^ hearts as indicated by wheat germ agglutinin staining (Figure 1D, E). No significant changes in cardiac fibrosis or collagen deposition were observed between PTP1B^+/+^::ꭤMHC^Cre/+^ and PTP1B^fl/fl^:: ꭤMHC^Cre/+^ mice from either sex, whether on ND or HFD (Figure S3). To more directly ascertain the PTP1B-associated effects on HFD-induced hypertrophy, we isolated individual CMs from both male and female PTP1B^+/+^::ꭤMHC^Cre/+^ and PTP1B^fl/fl^:: ꭤMHC^Cre/+^ hearts and found that the HFD PTP1B^+/+^::ꭤMHC^Cre/+^ cells were significantly enlarged in length, width, and overall area in response to HFD, as compared to ND (Figure 1F, S2D, Table 1, Table S3). In contrast, deletion of PTP1B showed a normal CM length, width, and overall area in response to HFD, similar in appearance to CMs isolated from mice fed a ND.

**Table 1.**
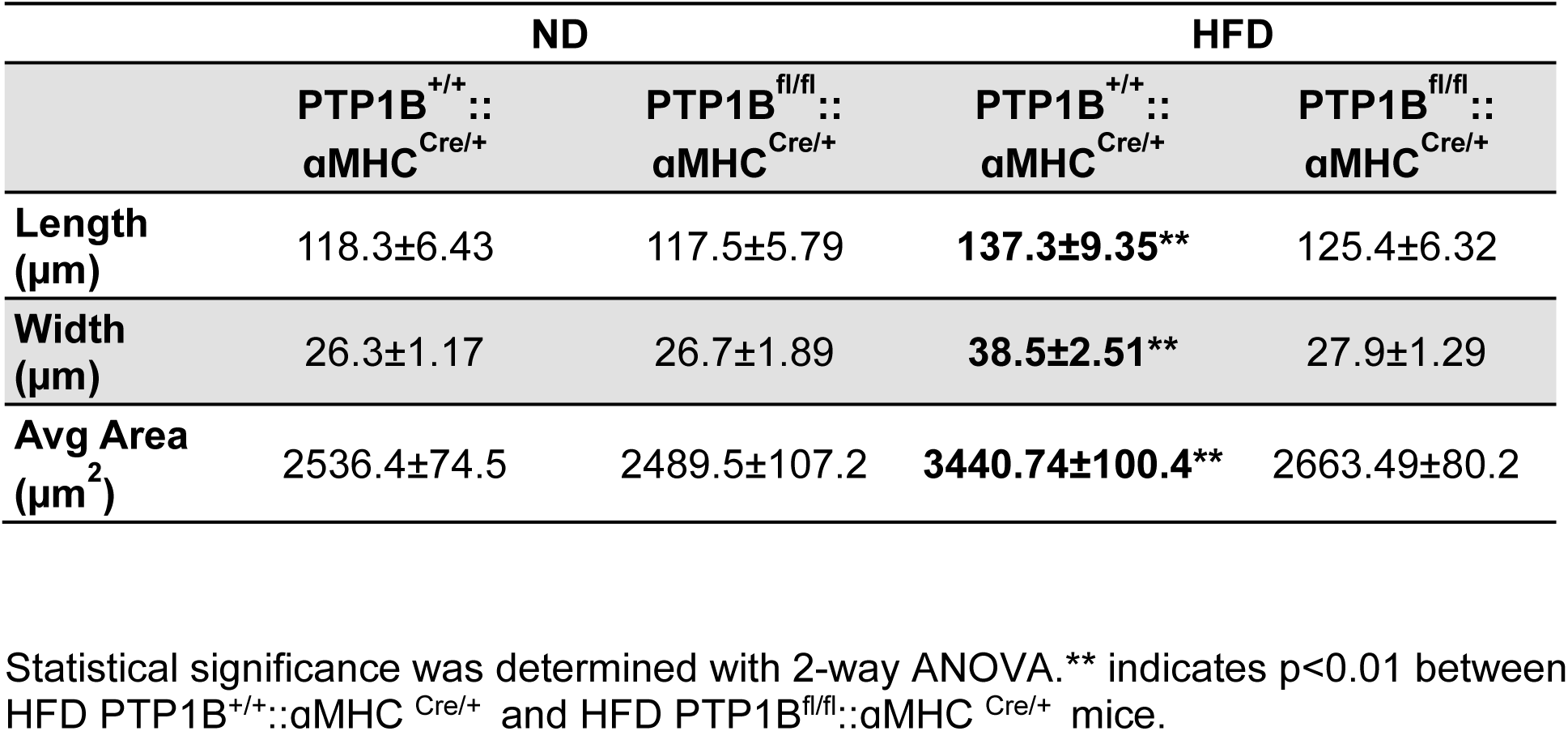
Measurements of isolated cardiomyocytes from male.

|  | ND |  | HFD |  |
| --- | --- | --- | --- | --- |
|  | PTP1B<sup>+/+</sup>::<br>αMHC<sup>Cre/+</sup> | PTP1B<sup>fl/fl</sup>::<br>αMHC<sup>Cre/+</sup> | PTP1B<sup>+/+</sup>::<br>αMHC<sup>Cre/+</sup> | PTP1B<sup>fl/fl</sup>::<br>αMHC<sup>Cre/+</sup> |
| **Length**<br>**(μm)** | 118.3±6.43 | 117.5±5.79 | **137.3±9.35\*\*** | 125.4±6.32 |
| **Width**<br>**(μm)** | 26.3±1.17 | 26.7±1.89 | **38.5±2.51\*\*** | 27.9±1.29 |
| **Avg Area**<br>**(μm<sup>2</sup>)** | 2536.4±74.5 | 2489.5±107.2 | **3440.74±100.4\*\*** | 2663.49±80.2 |

Statistical significance was determined with 2-way ANOVA. \*\* indicates p<0.01 between HFD PTP1B<sup>+/+</sup>::αMHC<sup>Cre/+</sup> and HFD PTP1B<sup>fl/fl</sup>::αMHC<sup>Cre/+</sup> mice.

Cardiac dimension and function were assessed by echocardiography in PTP1B^+/+^::ꭤMHC^Cre/+^ and PTP1B^fl/fl^::ꭤMHC^Cre/+^ mice following 8, 10 or 12 weeks of ND or HFD. While no functional echocardiographic parameters were affected in the females (Figure S4A-C, Table S4), we observed a significant protection in left ventricular posterior wall thickness (LVPW) and left ventricular chamber dimension in male PTP1B^fl/fl^::ꭤMHC^Cre/+^ hearts at 10 weeks and 12 weeks of HFD, respectively (Figure 3A C, Table 2). To determine if functional differences could be demonstrated at later time points in the females, we continued HFD treatment for an additional 8 weeks, to 20 weeks as described earlier (Fig S4B-C). Again, even at this later time point, we did not observe significant functional differences in the females in response to HFD.

**Table 2.**
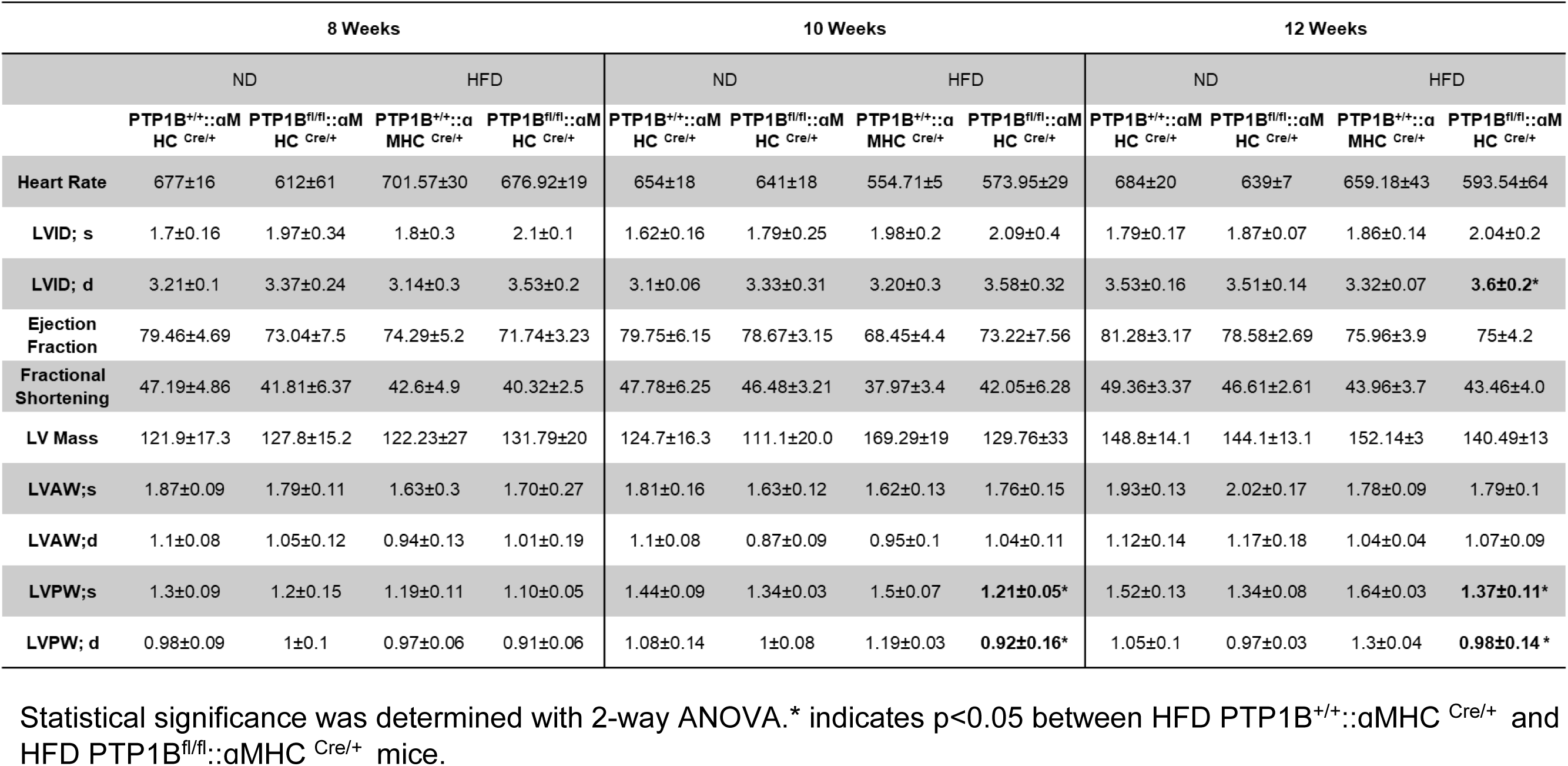
Echocardiographic analyses of male mice treated with ND or HFD for 8-12 weeks.

Assessment of fetal gene mRNA expression revealed a significant increase in atrial natriuretic factor (*ANF*) and a switch from αMHC (*MYH6*) to the β-myosin heavy chain (*MYH7*) in PTP1B^+/+^::ꭤMHC^Cre/+^, but not in PTP1B^fl/fl^:: ꭤMHC^Cre/+^HFD-induced CMs (Figure 2D). In contrast, we did not observe any significant changes in fetal gene expression in females for either control or PTP1B^fl/fl^:: ꭤMHC^Cre/+^ CMs in response to HFD (Figure S4D). Taken together, these results show that the deletion of PTP1B in CMs can attenuate cardiac hypertrophy in response to HFD, with a greater phenotype observed in males than females.

**Figure 2.**
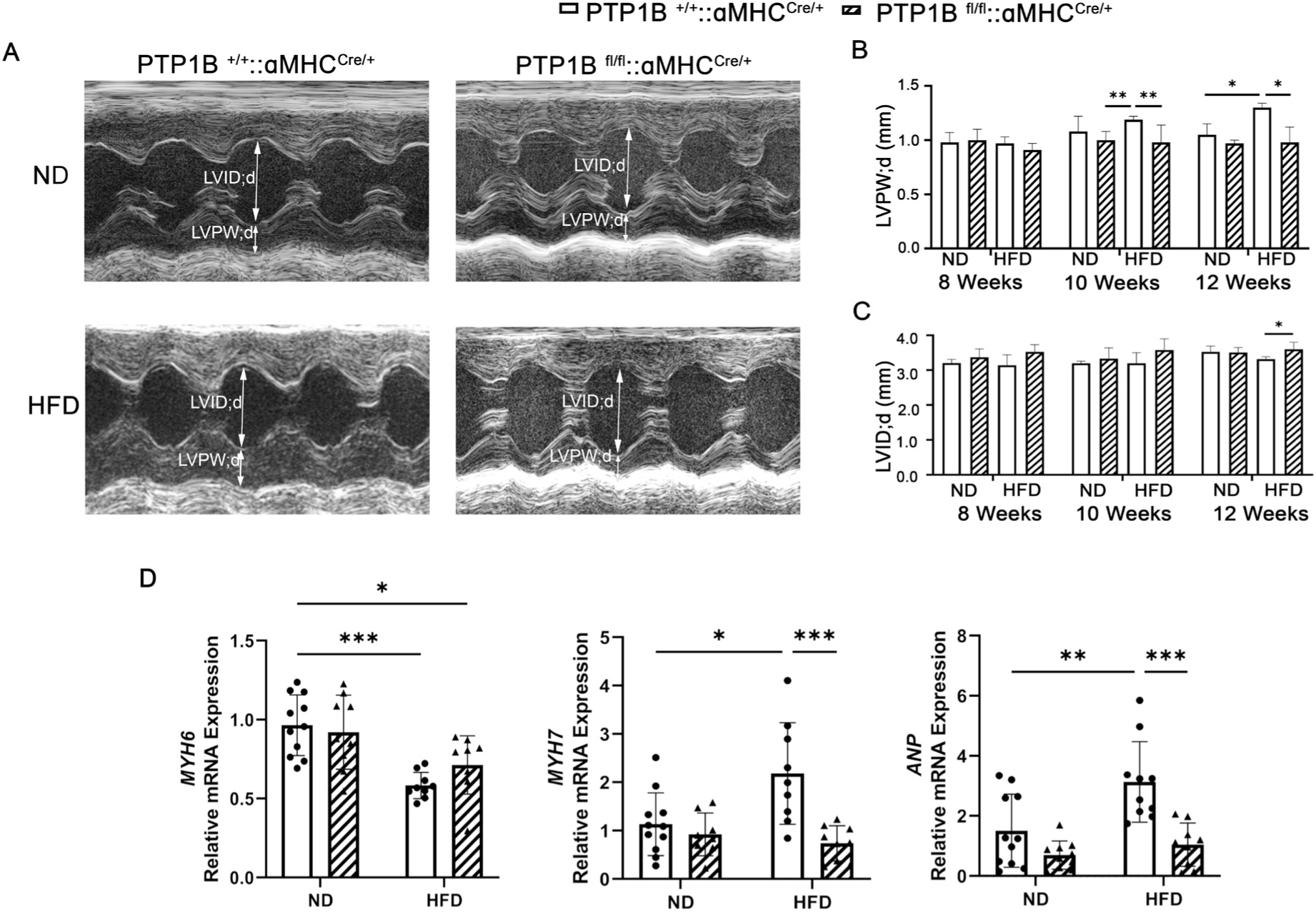
Cardiomyocyte-specific PTP1B deletion preserves cardiac structure in response to HFD. **A**. Representative echocardiography of PTP1B^+/+^::ꭤMHC ^Cre/+^ and PTP1B^fl/fl^::ꭤMHC ^Cre/+^ male mice fed either a ND or a HFD for up to 12 weeks. A two-headed arrow indicates left ventricular chamber size or left ventricular wall thickness. LVIDd= left ventricular internal diameter in end diastole. LVPWd=left ventricular posterior wall thickness in diastole. Quantification of **B.** LVPWd and **C.** LVIDd (n=10-15/group). **D.** Real-time q-PCR analysis of hypertrophy-related gene expression (*MYH6, MYH7, and ANP*) in CM mRNA isolated from either PTP1B^+/+^::ꭤMHC ^Cre/+^ or PTP1B^fl/fl^::ꭤMHC ^Cre/+^ male mice maintained on ND or HFD for 10 weeks. The ratio of ΔΔCT was analyzed using *18S* and eukaryotic elongation factor-1 (*Eef1*) mRNA as housekeeping controls, and HFD results were compared to ND from the same genotype. Samples were obtained from 6-7 mice per group and each sample was assessed in technical triplicate. Data in graphs represent mean ± SEM; *p<0.05, ** p<0.01, ***p<0.001, and all p values were derived from ANOVA with Bonferroni post-test when ANOVA was significant.

**Figure 3.**
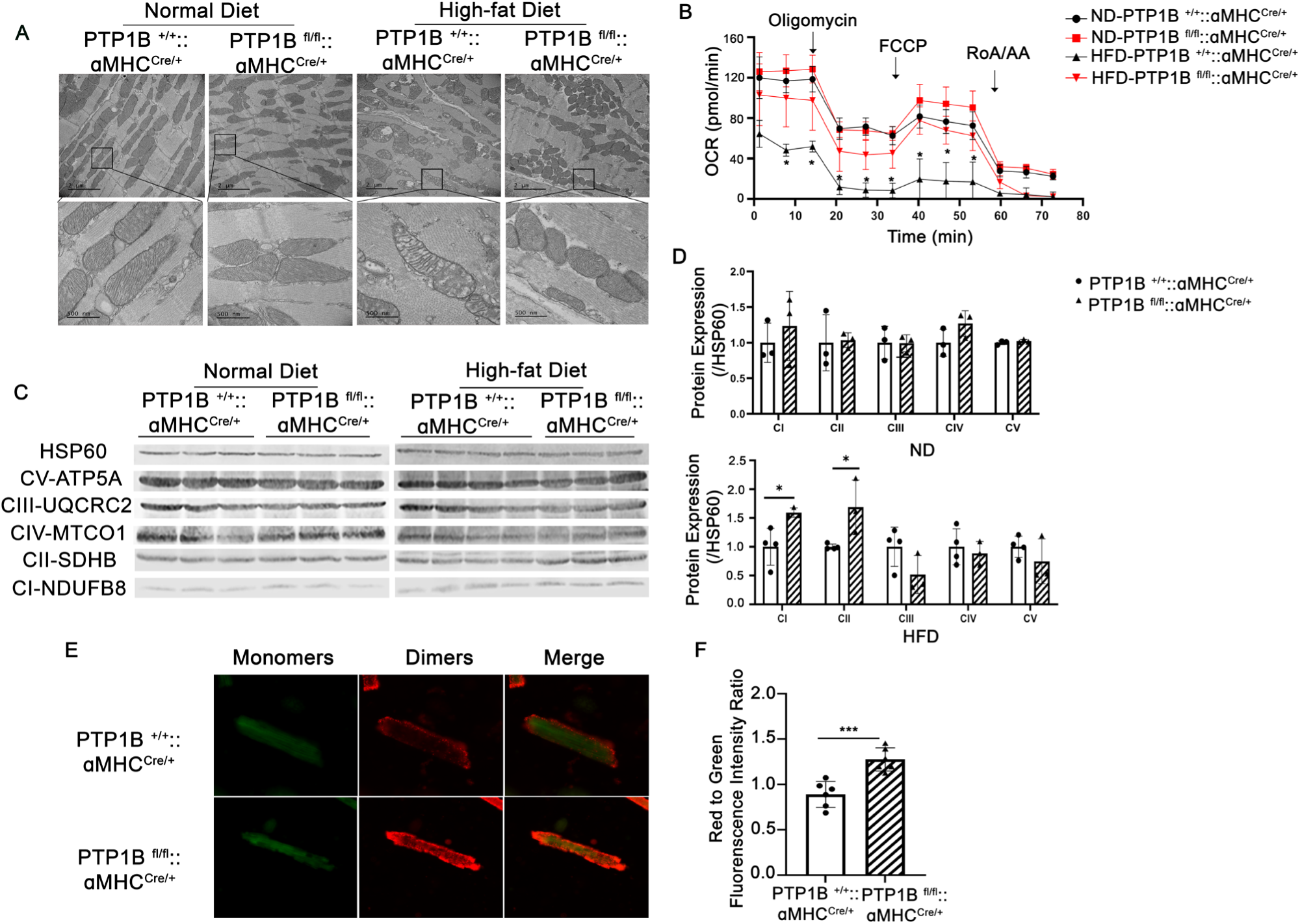
Cardiomyocyte-specific deletion of PTP1B preserves mitochondrial structure and energetics in presence of HFD. **A.** Representative TEM images of cardiac mitochondria from male PTP1B^+/+^::ꭤMHC ^Cre/+^ and PTP1B^fl/fl^::ꭤMHC ^Cre/+^ mice fed a ND or a HFD for 10 weeks. Scale bar= 2 µm (upper panel), 500 µm (lower panel). **B**. Oxygen consumption rates (OCR) measured in mitochondria isolated from PTP1B^+/+^::ꭤMHC ^Cre/+^ and PTP1B^fl/fl^::ꭤMHC ^Cre/+^ mice maintained under ND or HFD for 10 weeks. Measurements were done using a Seahorse analyzer after the sequential addition of oligomycin (Oligo, complex V inhibitor), FCCP (a protonophore), and antimycin A (AntiA, complex III inhibitor) to analyze ATP-linked respiration, proton leak respiration, maximal respiratory capacity, mitochondrial reserve capacity, and non-mitochondrial respiration. Data represent mean ± SEM, *p<0.05 using two-way ANOVA with Bonferroni post-hoc correction. **C**. Western blot and the **D.** quantitative assessment of OXPHOS mitochondrial complexes garnered from male PTP1B^+/+^::ꭤMHC ^Cre/+^ and PTP1B^fl/fl^::ꭤMHC ^Cre/+^ hearts on HFD for 10 weeks. Immunoblot utilized an antibody cocktail that recognizes the five mitochondrial oxidative phosphorylation complexes. *p<0.05 using one-way ANOVA. **E**. Representative image of JC-1 fluorescence in adult cardiomyocytes isolated from male PTP1B^+/+^::ꭤMHC ^Cre/+^ or PTP1B^fl/fl^::ꭤMHC ^Cre/+^ mice fed HFD for 10 weeks. Red fluorescence represents the JC-I mitochondrial aggregate, whereas green fluorescence indicates monomeric JC-1 (pathological). Scale bar=50 µm. **F.** Quantificative assessment of the red to green fluorescence intensity ratio, indicating changes in mitochondrial membrane potential (n= 6 mice per group). *p<0.05 is considered statistically significant after one-way ANOVA.

### Cardiomyocyte-specific deletion of PTP1B mitigates HFD-induced mitochondrial dysfunction

Mitochondrial dysfunction is a pathogenic hallmark of obesity-induced cardiomyopathy^51, 52^. Therefore, we investigated the effect of CM-specific PTP1B deletion on cardiac mitochondrial morphology and function. TEM showed no difference in mitochondrial ultrastructure between the PTP1B^+/+^::ꭤMHC^Cre/+^ and PTP1B^fl/fl^::ꭤMHC^Cre/+^ hearts under conditions of ND (Figure 3A). In contrast, we observed a profound disruption in mitochondrial ultrastructure, in which a subset of interfibrillar mitochondria was swollen with disorganized and reduced cristae density in the HFD-fed PTP1B^+/+^::ꭤMHC^Cre/+^ hearts. Remarkably, these changes were not observed in HFD-fed PTP1B^fl/fl^::ꭤMHC^Cre/+^ hearts, where mitochondrial integrity appeared normal and preserved (Figure 3A).

Next, we examined the effect of CM-specific PTP1B deletion on the respiratory activity of mitochondria using a Seahorse analyzer. Under conditions of ND, basal mitochondrial oxygen consumption rates (OCR) were similar between PTP1B^+/+^::ꭤMHC^Cre/+^ and PTP1B^fl/fl^::ꭤMHC^Cre/+^ mitochondria isolated from both male and female mouse hearts (Figure 3B). Moreover, OCR tracked similarly between ND-fed PTP1B^+/+^::ꭤMHC^Cre/+^ and PTP1B^fl/fl^::ꭤMHC^Cre/+^ mitochondria in the presence of the ATP synthase inhibitor oligomycin, the mitochondrial oxidative phosphorylation uncoupler FCCP, and following treatment with rotenone and antimycin (Figure 3B, S5A). In contrast, HFD-fed PTP1B^+/+^::ꭤMHC^Cre/+^, but not PTP1B^fl/fl^::ꭤMHC^Cre/+^ isolated mitochondria, showed significant impairment of OCR, with reduced overall baseline levels of respiration, suggesting deletion of PTP1B in CMs protects against HFD-induced mitochondrial dysfunction in both male and female mouse hearts (Figure 3B).

Next, we examined the protein levels of electron transport chain (ETC) complexes. We observed comparable levels of all OXPHOS complexes in ND-fed PTP1B^+/+^::ꭤMHC^Cre/+^ and PTP1B^fl/fl^::ꭤMHC^Cre/+^ hearts, in both males and females (Figure 3C,3D, S5A-B). In response to HFD, however, expression of NADH: ubiquinone oxidoreductase subunit B8 (NDUFB8; Complex I) and succinate dehydrogenase subunit b (SDHB; Complex II) were significantly increased in PTP1B^fl/fl^::ꭤMHC^Cre/+^ mouse hearts, relative to control HFD hearts, raising the possibility of increased flux through complex I (Figure 3C, 3D, S5A-B).

To evaluate the mitochondrial membrane potential, we used JC-1 to stain CMs isolated from HFD-fed PTP1B^+/+^::ꭤMHC^Cre/+^ and PTP1B^fl/fl^::ꭤMHC^Cre/+^ mice. Cells with high mitochondrial membrane potential (Δψ) promote the formation of red fluorescent JC-1 aggregates, while cells with low Δψ exhibit green fluorescence^53^. We found that PTP1B^fl/fl^::ꭤMHC ^Cre/+^ CMs have highly active and hyperpolarized mitochondria, as compared to mitochondria from isolated CMs of PTP1B^+/+^::ꭤMHC^Cre/+^ mice, confirming that the depletion of PTP1B in CMs prevents HFD-induced mitochondrial dysfunction (Figure 3E, 3F). Together, these results indicate that CM-specific deletion of PTP1B preserves mitochondrial function and structural integrity following HFD.

### Cardiac-specific deletion of PTP1B shifts cardiac metabolism from glycolysis to fatty acid oxidation

To better characterize the metabolic phenotype of CM-specific deletion of PTP1B, we analyzed the cardiac tissue metabolome of HFD-fed PTP1B^+/+^::ꭤMHC^Cre/+^ and PTP1B^fl/fl^::ꭤMHC^Cre/+^ mice. Our resulting dataset identified a total of 741 metabolites, 86 of which were significantly different between control and PTP1B^fl/fl^::ꭤMHC^Cre/+^ hearts (Summarized in Table S5). Indeed, many of these metabolites affect critical metabolic signaling pathways (Tables 3-7). Specifically, levels of metabolites central to glycolysis, such as glucose 6-phosphate, fructose 6-phosphate, fructose 1,6-bisphosphate, were decreased in both male and female PTP1B^fl/fl^::ꭤMHC^Cre/+^ mouse hearts, as compared to controls (Table 3). Moreover, HFD-fed PTP1B^fl/fl^::ꭤMHC^Cre/+^ mouse hearts exhibited decreased TCA cycle intermediates (Table 4). We next measured the activity of pyruvate dehydrogenase (PDH), an enzyme that catalyzes pyruvate into acetyl-CoA, the rate limiting step for glucose oxidation^54–56^ ; increased phosphorylation of PDH is indicative of decreased PDH activity. Our data show that PDH phosphorylation was increased in HFD fed PTP1B^fl/fl^::ꭤMHC^Cre/+^ mouse hearts, as compared to controls (Figure 4A, 4B, S6A, S6B).

**Figure 4.**
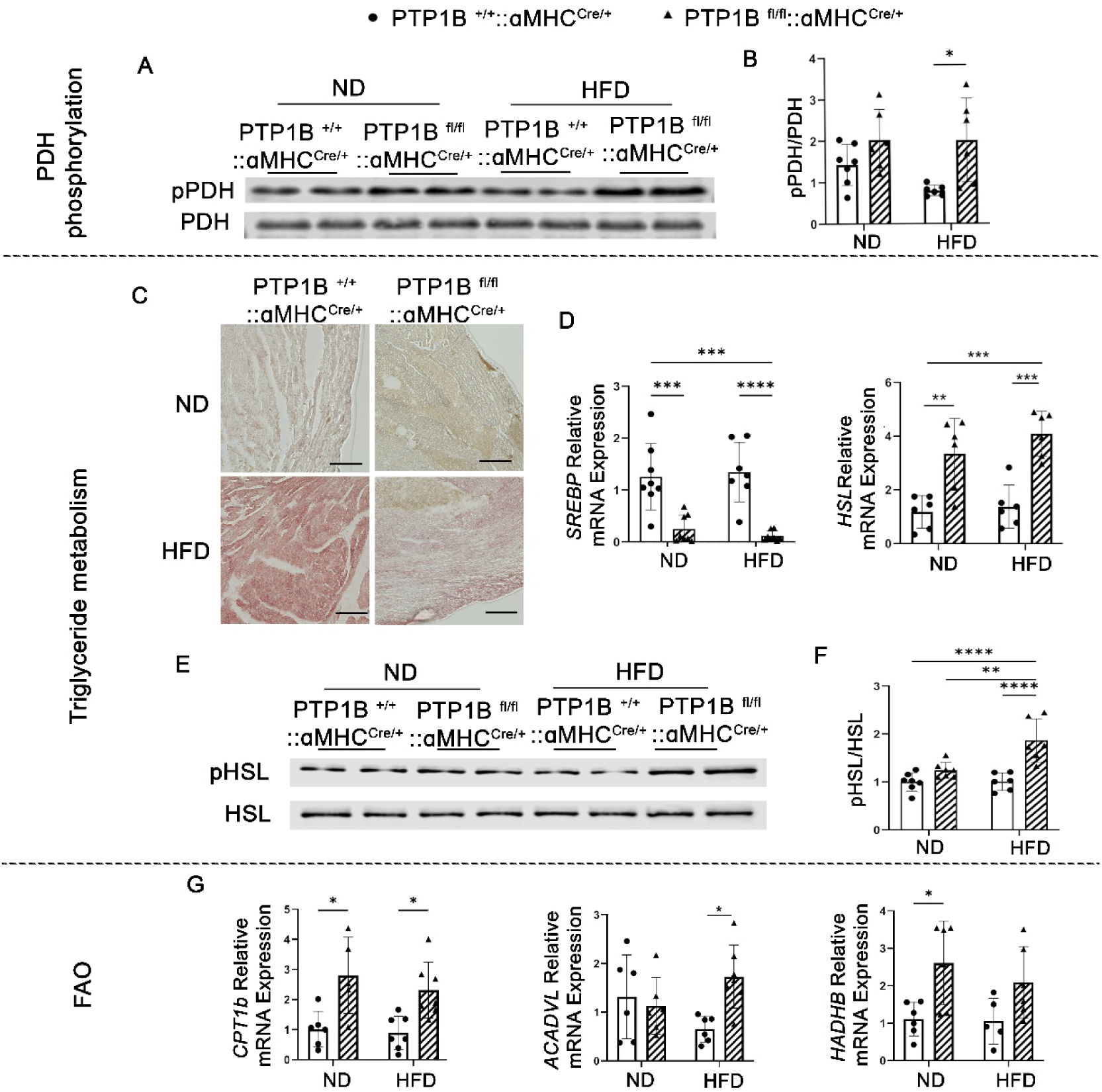
Triglyceride mobilization and fatty acid oxidation are increased in PTP1B^fl/fl^::ꭤMHC ^Cre/+^ hearts. **A.** Representaative western blot of heart lysates immunoblotted with phospho-PDH (Ser 293) or PDH antibodies. **B.** Quantification of pPDH/total PDH ratios. N=6-7 mice/group. **C.** Oil Red O staining lipid droplets in frozen male heart sections from both PTP1B^+/+^::ꭤMHC^Cre/+^ and PTP1B^fl/fl^::ꭤMHC^Cre/+^ mice fed either ND or HFD for 10 weeks. Scale bar= 50 µm. **D**. Gene mRNA expression analysis (real-time PCR) of lipogenic transcription factors *SREBP* and *HSL* in CMs isolated from PTP1B^+/+^::ꭤMHC^Cre/+^ and PTP1B^fl/fl^::ꭤMHC^Cre/+^. The ratio of ΔΔCT was analyzed using *18S* and eukaryotic elongation factor-1 (*Eef1*) mRNA as housekeeping controls, and HFD results were compared to ND from the same genotype. Samples were obtained from 7-9 mice per group and each sample was assessed in technical triplicate. Data in graphs represent mean ± SEM; *p<0.05, ** p<0.01, ***p<0.001, and all p values were derived from ANOVA with Bonferroni post-test when ANOVA was significant. **E**. Representative western blotting analysis and **F.** quantitative analysis of HSL protein expression in heart lysates isolated from PTP1B^+/+^::ꭤMHC^Cre/+^ or PTP1B^fl/fl^::ꭤMHC^Cre/+^ mice fed HFD for 10-weeks. N=6-7 mice/group. **G**. Gene expression of fatty acid oxidation genes *CPT1b*, *ACADVL*, and *HADHB* in CMs isolated from PTP1B^+/+^::ꭤMHC^Cre/+^ or PTP1B^fl/fl^::ꭤMHC^Cre/+^. The ratio of ΔΔCT was analyzed using *18S* and eukaryotic elongation factor-1 (*Eef1*) mRNA as housekeeping controls, and HFD results were compared to ND from the same genotype. Samples were obtained from 7-9 mice per group and each sample was assessed in technical triplicate. Data in graphs represent mean ± SEM; *p<0.05, ** p<0.01, ***p<0.001, and all p values were derived from ANOVA with Bonferroni post-test when ANOVA was significant. Statistical significance was determined with 2-way ANOVA. *p<0.05, ** p<0.01, ***p<0.001.

**Table 3.**
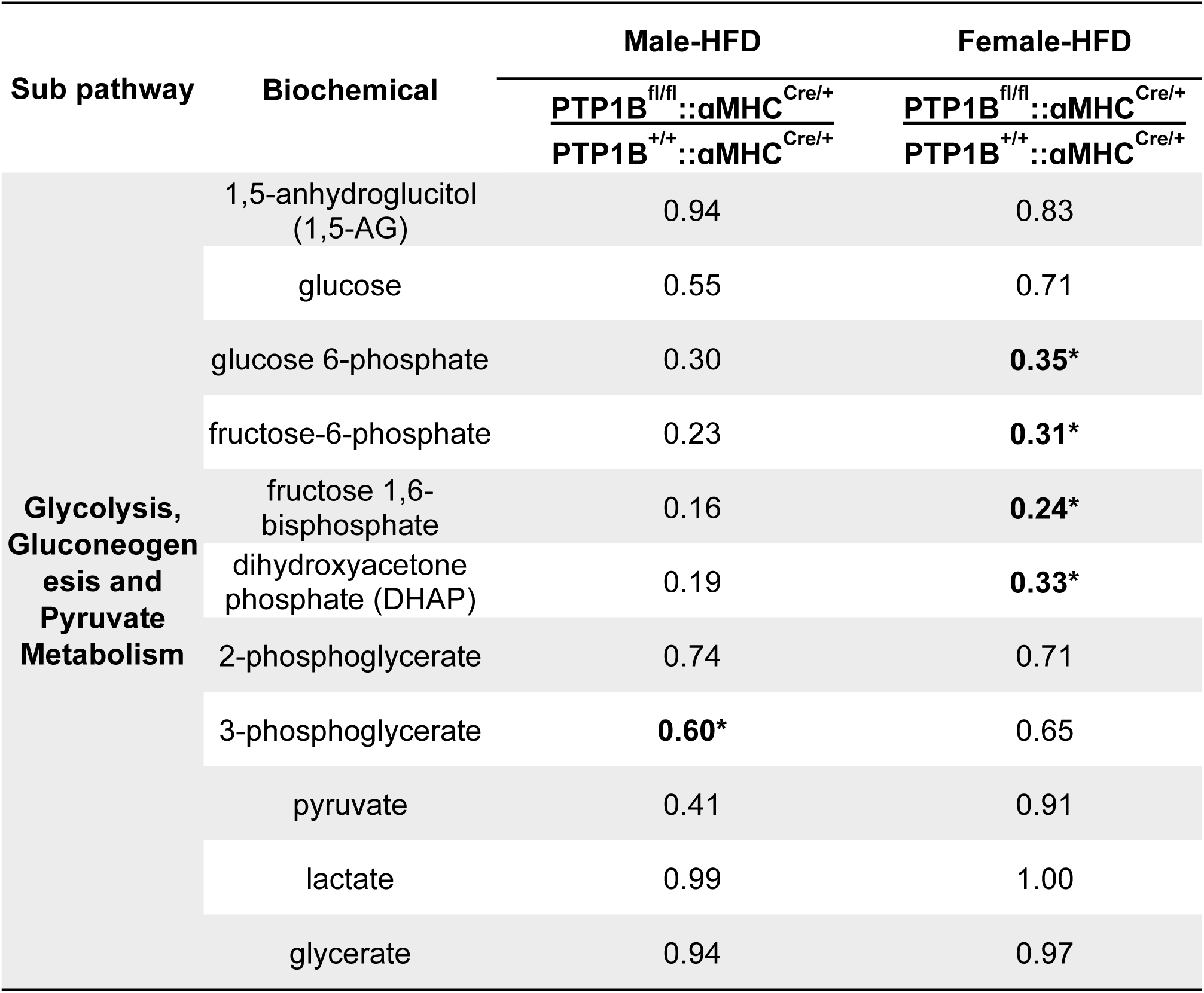
Relative ratios ofglucose metabolism intermediates between PTP1B^+/+^::ꭤMHC^Cre/+^ and PTP1B^fl/fl^::ꭤMHC^Cre/+^ mouse hearts.

**Table 4.** Relative ratios of TCA cycle intermediates between PTP1B^+/+^::ꭤMHC^Cre/+^ and PTP1B^fl/fl^::ꭤMHC^Cre/+^ mouse hearts.

| Sub pathway | Biochemical | Male-HFD | Female-HFD |
| --- | --- | --- | --- |
| | | $\frac{\text{PTP1B}^{\text{fl/fl}} :: \alpha\text{MHC}^{\text{Cre/+}}}{\text{PTP1B}^{+/+} :: \alpha\text{MHC}^{\text{Cre/+}}}$ | $\frac{\text{PTP1B}^{\text{fl/fl}} :: \alpha\text{MHC}^{\text{Cre/+}}}{\text{PTP1B}^{+/+} :: \alpha\text{MHC}^{\text{Cre/+}}}$ |
| TCA Cycle | succinylcarnitine (C4-DC) | **0.6\*** | 1.11 |
|  | acetyl CoA | 0.89 | **0.78\*** |
|  | glutamate | 0.96 | **0.78\*** |
|  | glutamine | 0.9 | 0.92 |
|  | glutamate, gamma-methyl ester | 1.02 | 0.84* |

Next, we asked if the utilization of fat is altered in PTP1B^fl/fl^::ꭤMHC^Cre/+^ hearts. Oil red O staining showed that normal diet does not lead to lipid accumulation in either male or female PTP1B^+/+^::ꭤMHC^Cre/+^ or PTP1B^fl/fl^::ꭤMHC^Cre/+^ mouse hearts (Figure 4C, upper left and right panels). In contrast, high-fat diet significantly promoted lipid accumulation in PTP1B^+/+^::ꭤMHC^Cre/+^ (Figure 4C and S6C, lower left panels), but not in PTP1B^fl/fl^::ꭤMHC ^Cre/+^ hearts (Figure 4C and S6C, lower right panels). To evaluate the effects of PTP1B on myocardial lipid metabolism, we measured the expression of lipogenic genes by quantitative real-time polymerase chain reaction (qRT-PCR). Our results reveal that while high-fat diet significantly increased the cardiac lipogenic gene *SREBP1c* in PTP1B^+/+^::ꭤMHC^Cre/+^, the expression of this gene was significantly reduced in PTP1B^fl/fl^::ꭤMHC^Cre/+^ hearts (Figure 4D, S6D). In addition, expression levels and phosphorylation of the key enzyme (HSL), which mediates triglyceride hydrolysis, was increased in PTP1B^fl/fl^::ꭤMHC^Cre/+^ hearts, relative to controls (Figure 4D-F, S6D-F), which paralleled reduced lipid droplet accumulation in cardiac tissues of these mice. FAO metabolites, acylcarnitines (AC), were also significantly increased in both male and female PTP1B^fl/fl^::ꭤMHC^Cre/+^ mouse hearts, as compared to controls (Table 5). ACs arise from the conjugations of acyl-coenzyme A with carnitine for the transport of long-chain fatty acids across the inner mitochondrial membrane for β-oxidation. To evaluatethese findings, we measured key metabolic enzymes by qRT-PCR. We found that CM-specific deletion of PTP1B increases expression levels of the mitochondrial long chain fatty acyl importer, *CPT1b* in response to both ND and HFD (Figure 4G, S6G). Acyl-CoA dehydrogenase very long chain (*ACADVL*), which encodes the protein that catalyzes the first step of the mitochondrial beta oxidation pathway, is also elevated in HFD-fed PTP1B^fl/fl^::ꭤMHC^Cre/+^ mice (Figure 4G, S6G). Similarly, the mRNA level of acetyl-CoA acyltransferase (*HADHB*) was also increased in PTP1B^fl/fl^::ꭤMHC^Cre/+^ mouse hearts in response to both ND and HFD (Figure 4G, S6G). Taken together, these data suggest that CM-specific PTP1B deletion could increase cardiac FAO while decreasing glucose utilization.

**Table 5.** Relative ratio of fatty acid metabolism intermediates in PTP1B^+/+^::ꭤMHC^Cre/+^ and PTP1B^fl/fl^::ꭤMHC^Cre/+^ mouse hearts.

| Sub pathway | Biochemical | Male-HFD | Female-HFD |
| --- | --- | --- | --- |
| | | $\frac{\text{PTP1B}^{\text{fl/fl}}::\alpha\text{MHC}^{\text{Cre/+}}}{\text{PTP1B}^{+/+}::\alpha\text{MHC}^{\text{Cre/+}}}$ | $\frac{\text{PTP1B}^{\text{fl/fl}}::\alpha\text{MHC}^{\text{Cre/+}}}{\text{PTP1B}^{+/+}::\alpha\text{MHC}^{\text{Cre/+}}}$ |
| **Acyl Carnitine** | cis-4-decenoylcarnitine (C10:1) | 1.58 | 1.14 |
|  | margaroylcarnitine (C17) | **2.47\*** | **1.68\*** |
|  | arachidoylcarnitine (C20) | **1.76\*** | **3.00\*** |
|  | erucoylcarnitine (C22:1)* | **3.09\*** | **3.34\*** |
|  | behenoylcarnitine (C22) | 1.06 | **4.26\*** |
|  | lignoceroylcarnitine (C24)* | 1.10 | **2.78\*** |
|  | nervonoylcarnitine (C24:1) | 1.19 | **3.57\*** |
|  | carnitine | **1.35** | 0.95 |

### Cardiomyocyte-specific deletion of PTP1B increases metabolic signaling

PTP1B plays an integral role in modulating multiple receptor tyrosine kinase signaling pathways, including downstream effectors such as PI3K/AKT, Ras/MAPK, and PKM2/AMPK (Figure S7)^23, 33, 36, 57^. For example, PTP1B is a known negative regulator of IR and VEGFR^58–60^. To begin to assess the molecular consequences of PTP1B deletion in the heart, we intraperitoneally injected PTP1B^+/+^::ꭤMHC^Cre/+^ or PTP1B^fl/fl^::ꭤMHC^Cre/+^ mice with saline (baseline) or insulin for 10 minutes and measured the effects on IR or VEGFR phosphorylation. While we did not observe significant differences at baseline, following insulin stimulation, hearts isolated from PTP1B^fl/fl^::ꭤMHC^Cre/+^ mice fed either ND or HFD exhibited hyper-activation of both IR and VEGFR, as compared to insulin-stimulated control mouse hearts (Figure 5A-C).

**Figure 5.**
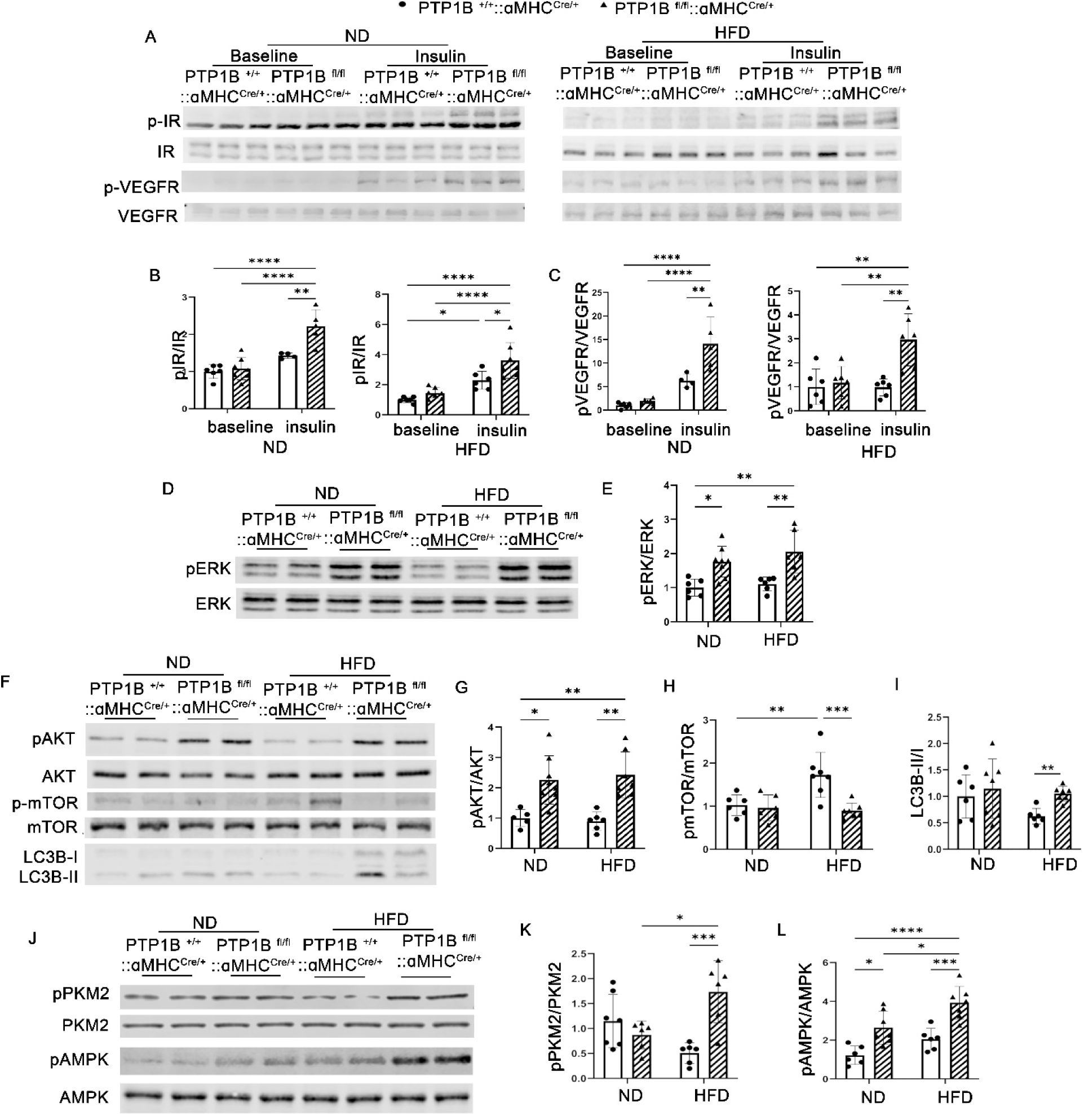
Loss of PTP1B in CMs induces activation of ERK, AKT, PKM2 and AMPK. **A.** Representative western blot of phospho and total insunlin receptor or VEGFR in cardiac extracts isolated from male PTP1B^+/+^::ꭤMHC^Cre/+^ or PTP1B^fl/fl^::ꭤMHC^Cre/+^ mice fed a ND or HFD injected with saline or insulin (10 mU/g i.p.). **B.** Quantificatative analyses of the ratio of IR phosphorylation (Tyr1162/1163) to total IR expression generated from three independent experiments, n=6-8 hearts/group. **C**. Quantificatative analyses of the ratio of VEGFR phosphorylation (Tyr1175) to total VEGFR expression generated from three independent experiments, n=6-8 hearts/group. **D**. Representative western blot of phospho and total ERK in cardiac extracts isolated from male PTP1B^+/+^::ꭤMHC^Cre/+^ or PTP1B^fl/fl^::ꭤMHC^Cre/+^ mice fed a ND or HFD. **E.** Quantificatative analyses of the ratio of ERK phosphorylation to total ERK expression generated from three independent experiments, n=6-8 hearts/group. **F-I.** Representative western blots and quantitative analysis of total and phospho-AKT (Ser 473), total and phospho-mTOR (Ser 248), and autophagy markers, LC3I and LC3II in cardiac extracts isolated from male PTP1B^+/+^::ꭤMHC^Cre/+^ or PTP1B^fl/fl^::ꭤMHC^Cre/+^ mice fed ND or HFD. n=6-8 hearts/group. . **J-L**. Representative western blots and quantitative analyis of heart lysates immunoblotted with total and phospho-PKM2 (Tyr105), as well as total and phospho-AMPK (Thr 172). n=6-8 hearts/group.

To further understand the effects of cardiac-specific PTP1B deletion on these pathways, we measured changes in ERK activation, a downstream effector of VEGFR. Hearts isolated from both male and female PTP1B^fl/fl^::ꭤMHC^Cre/+^ mice fed either ND or HFD showed significant increases in phosphorylated ERK (Figure 5D, 5E, S7A, S7B). We next assessed the effects of PTP1B deletion on the PI3K/AKT signaling pathway. Here too, we found that both male and female PTP1B^fl/fl^::ꭤMHC^Cre/+^ mouse hearts had significantly increased levels of AKT phosphorylation in response to both ND and HFD, as compared to control (Figure 5F, 5G, S7C, S7D). AKT-dependent enhancement of protein synthesis is mediated in part by activation of mammalian target of rapamycin (mTOR). However, though we observed an increase in mTOR phosphorylation in HFD-fed PTP1B^+/+^::ꭤMHC^Cre/+^ mouse hearts, PTP1B^fl/fl^::ꭤMHC^Cre/+^ hearts surprisingly did not reveal altered mTOR activation (Figure 5F, 5H, S7C, S7E). Concomitantly, HFD-fed PTP1B^fl/fl^::ꭤMHC^Cre/+^ hearts also had increased downstream autophagy, as compared to HFD-fed control mouse hearts, as demonstrated by increased LC3B-II-to LC3B-I ratios (Figure 5F, 5I, S7C, S7F). Together, these data suggest that while PTP1B affects both ERK and AKT activities, a possible parallel signaling pathway may be the driver that differentially modulates mTOR and autophagy in response to HFD in these CM-specific deleted PTP1B mice.

While HFD has been shown to induce mTOR activity in the liver and skeletal muscle, leading to impaired insulin signaling^61^, the fact that CM-specific deletion of PTP1B reduces mTOR activity in response to HFD suggests a cardio-protective role for PTP1B against HFD-induced insulin resistance. However, the mechanism for how this occurs remains unclear. Previously, PKM2, a rate-limiting glycolytic enzyme, was shown to be dephosphorylated and activated by PTP1B in both pancreatic cancer cells and in cultured adipocytes^33, 36^ Figure S7). To assess whether CM-specific PTP1B affects this pathway, we examined the phosphorylation status of PKM2 in mouse hearts isolated from either ND or HFD-fed control or PTP1B^fl/fl^::ꭤMHC^Cre/+^ mice. We found that PKM2 phosphorylation was significantly elevated only in HFD-fed PTP1B^fl/fl^::ꭤMHC^Cre/+^ hearts (Figure 5J, 5K, S7G, S7H), suggesting decreased PKM2 activity. Concomitantly, because PKM2 normally functions to inhibit AMPK activity, we found that AMPK activity was significantly increased in PTP1B^fl/fl^::ꭤMHC^Cre/+^ hearts, even under ND conditions (Figure 5J, 5L, S7G, S7I). Together, these results suggest that the absence of PTP1B in CMs improves insulin resistance and HFD-associated cardiomyopathy by increasing AMPK in the heart.

### PTP1B differentially regulates NAD^+^ to modulate cardiac metabolic functions in response to HFD

Our data suggest that HFD-fed PTP1B^fl/fl^::ꭤMHC^Cre/+^ mice exhibit a significant alteration in their metabolic signaling and metabolic profiles, as compared to HFD-fed control mice. Moreover, AMPK has been reported to mediate NAD^+^ metabolism^62^. Therefore, we hypothesized that because of the effects we see on the PKM2/AMPK signaling axis, that CM-specific deletion of PTP1B protects against impaired HFD-induced cardiomyopathy through activation of the NAD^+^ biosynthesis pathway. To begin to assess this, we first measured changes in NAD^+^ metabolic intermediates. Interestingly, we observed increased levels of nicotinamide ribonucleotide (NMN) in male, but not female, PTP1B^fl/fl^::ꭤMHC^Cre/+^ hearts, as compared to controls (Table 6). However, both male and female PTP1B^fl/fl^::ꭤMHC^Cre/+^ mouse hearts had diminished nicotinamide (NAM) metabolites (Table 6), suggesting a significant role for this pathway in the regulation of PTP1B in the heart.

**Table 6.**
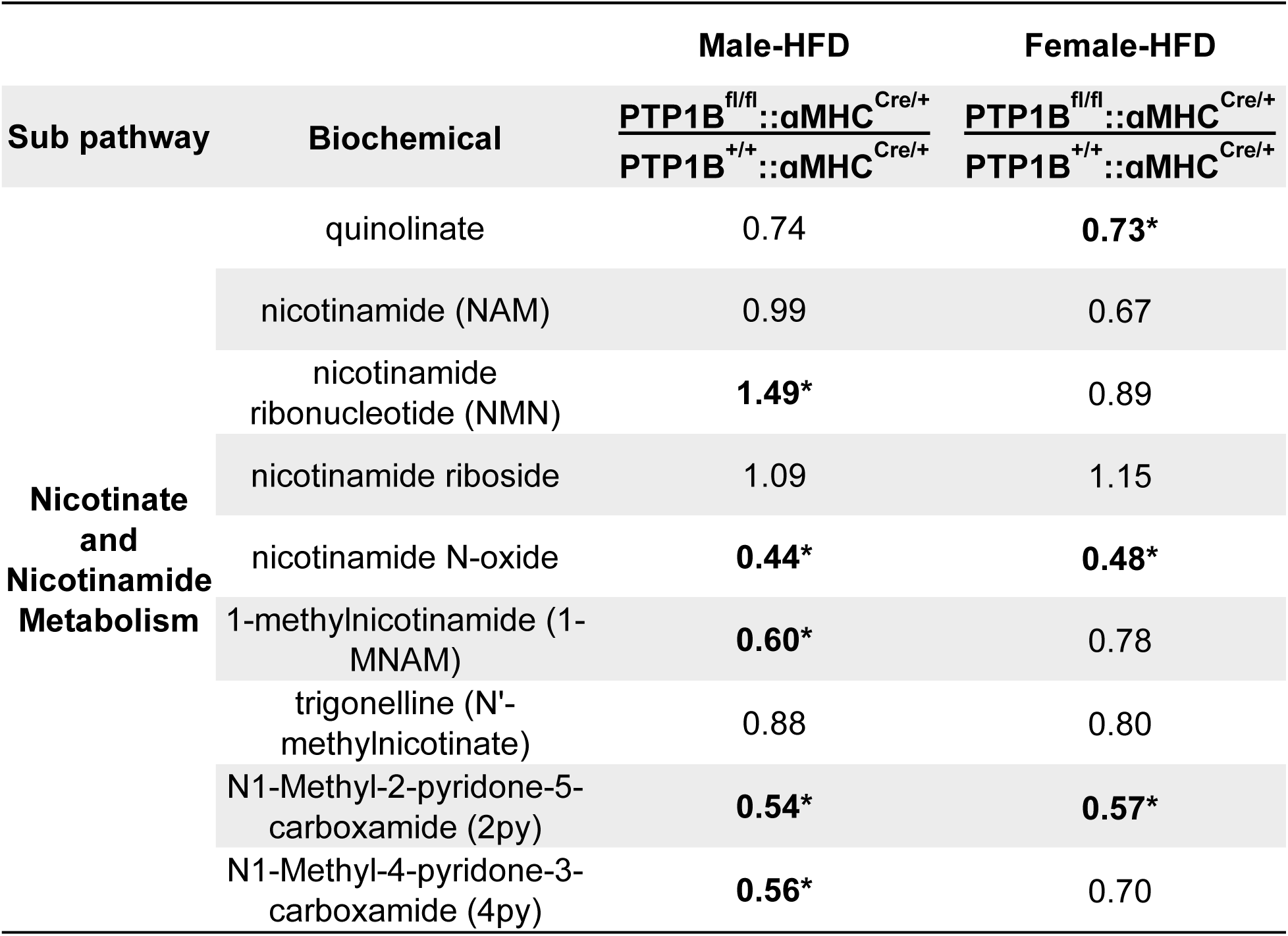
Relative ratios of NAD^+^ metabolism intermediates between PTP1B^+/+^::ꭤMHC^Cre/+^ and PTP1B^fl/fl^::ꭤMHC^Cre/+^ mouse hearts.

To validate this, we measured the levels of NAD^+^, NADH, NADP^+^, and NADPH directly in hearts of either PTP1B^+/+^::ꭤMHC^Cre/+^ or PTP1B^fl/fl^::ꭤMHC^Cre/+^. We found that all NAD^+^ intermediates were increased in CM-specific deleted PTP1B hearts, as compared to controls (Figure 6A). Next, we measured the transcriptional expression of genes related to NAD^+^ synthesis. Levels of *NAMPT*, *NMNAT*, *NAPRT*, and *NADSYN* were all significantly higher in hearts from both male and female PTP1B^fl/fl^::ꭤMHC^Cre/+^ mice, in response to both ND and HFD (Figure 6B). Moreover, the expression of NAMPT was also elevated in hearts isolated from PTP1B^fl/fl^::ꭤMHC^Cre/+^ mice (Figure 6C, 6D). Taken together, these results suggest that CM-specific deletion of PTP1B promotes NAD^+^ biosynthesis through the regulation of metabolic processes in cardiac mitochondria and regulation of metabolic signaling pathways regulated primarily by AMPK activity.

**Figure 6.**
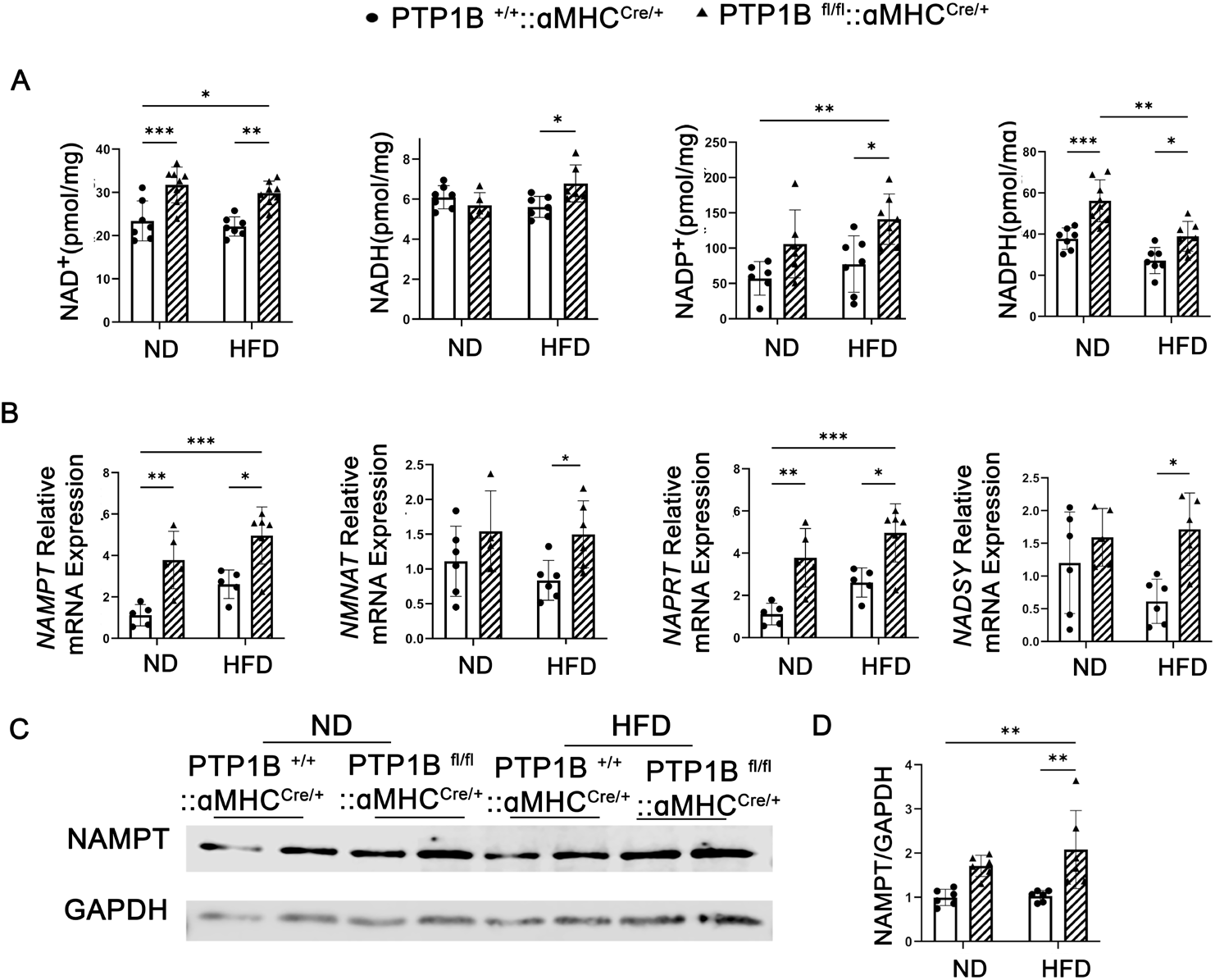
NAD^+^ metabolites are increasedin PTP1B^fl/fl^::ꭤMHC ^Cre/+^ mice. **A**. Levels of cardiac NAD^+^, NADH, NADP^+^ and NADPH were evaluated in male PTP1B^+/+^::ꭤMHC ^Cre/+^ and PTP1B^fl/fl^::ꭤMHC ^Cre/+^ mice after either ND or HFD for 10 weeks. **B.** NAD^+^ salvage synthesis pathway critical genes *NAMPT*, *NMNAT*, *NAPRT*, *and NADSYN* were measured by quantitative real-time PCR in CMs from PTP1B^+/+^::ꭤMHC ^Cre/+^ and PTP1B^fl/fl^::ꭤMHC ^Cre/+^ mice fed ND or HFD for 10-weeks. The ratio of ΔΔCT was analyzed using *18S* and eukaryotic elongation factor-1 (*Eef1*) mRNA as housekeeping controls, and HFD results were compared to ND from the same genotype. Samples were obtained from 7-9 mice per group and each sample was assessed in technical triplicate. Data in graphs represent mean ± SEM; *p<0.05, ** p<0.01, ***p<0.001, and all p values were derived from ANOVA with Bonferroni post-test when ANOVA was significant. Statistical significance was determined with 2-way ANOVA. *p<0.05, ** p<0.01, ***p<0.001. **C-D**. Representative western blots and quantitative analyis of heart lysates immunoblotted with NAMPT and GAPDH (control for loading). N=6-8 hearts/group).. Statistical significance was determined with 2-way ANOVA. *p<0.05, ** p<0.01, ***p<0.001.

## Discussion

Using a HFD-induced cardiomyopathy mouse model, we made a series of novel mechanistic observations: (1) CM-specific deletion of PTP1B in mice ameliorates HFD induced cardiomyopathy (ie, hypertrophy) and diminishes cardiac steatosis; (2) CM specific deletion of PTP1B activates AKT and ERK signaling, but decreases mTOR phosphorylation; (3) the suppression of mTOR is regulated by a novel PTP1B-PKM2 AMPK axis in the heart, increasing autophagic activity; (4) deletion of PTP1B in CMs alters lipid metabolism and mitochondrial function in response to obesity-induced cardiomyopathy; (5) CM-specific PTP1B deletion may promote more complete cardiac FAO while decreasing glycolysis in response to HFD; (6) CM-specific deletion of PTP1B promotes NAD^+^ biosynthesis through AMPK-mediated metabolic signaling pathways (Figure 7, S7). Taken together, our data indicate that CM-specific deletion of PTP1B protects the heart against development of HFD-induced cardiomyopathy through its direct regulation of cardiac metabolic signaling.

**Figure 7.**
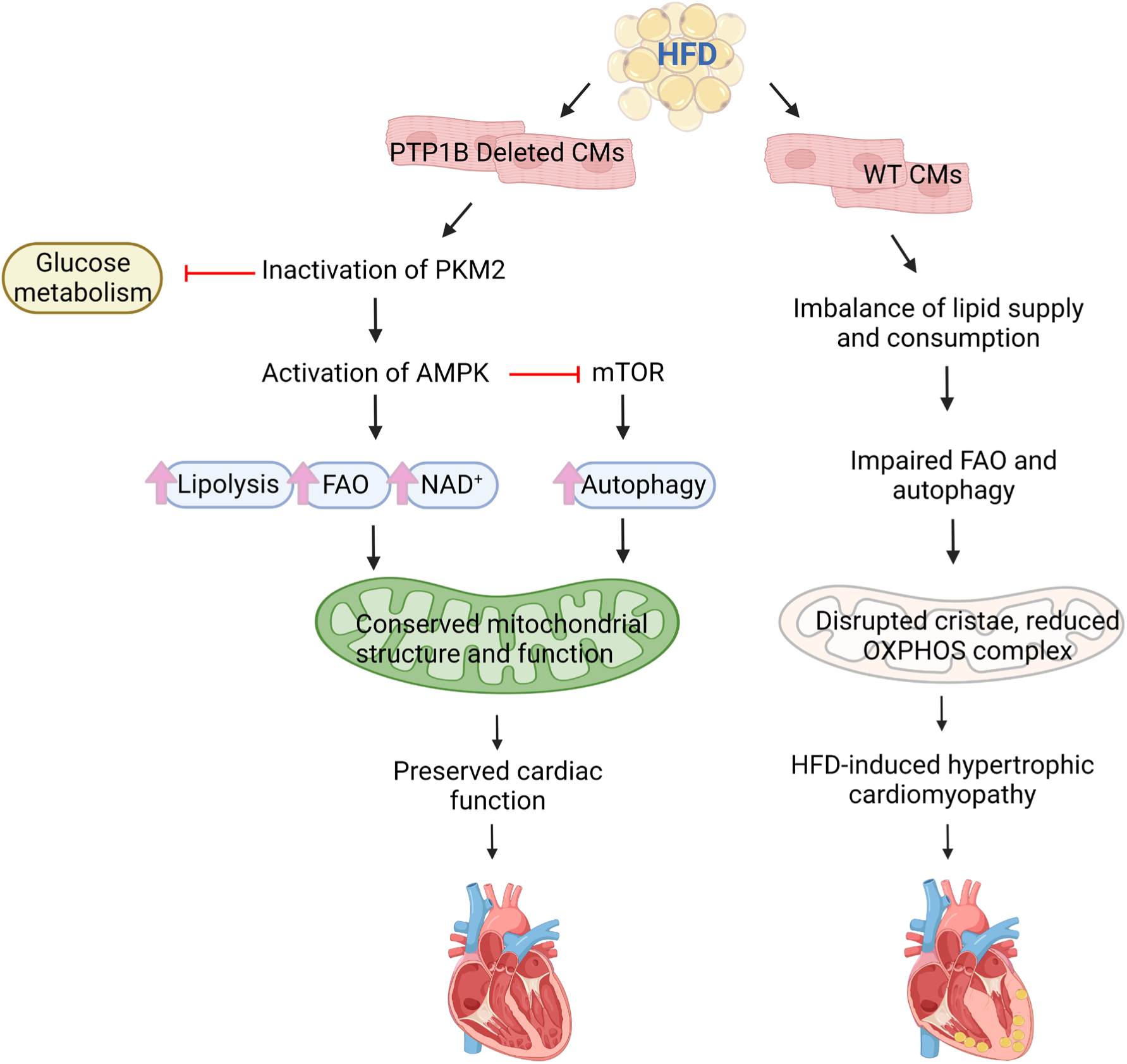
Schematic summary for the metabolic role of PTP1B in the heart. PTP1B functions as a cardio-metabolic switch in response to high-fat diet induced cardiomyopathy. Cardiomyocyte-specific deletion of PTP1B inactivates PKM2 and activates AMPK, leading to increased autophagy, fatty acid oxidation and NAD^+^ synthesis, thereby protecting the heart against HFD-induced cardiomyopathy.

Clinically, studies have found increased PTP1B activity in patients with systolic dysfunction ^43^. Conversely, germline deletion of PTP1B in mice improves cardiac output without affecting infarct size^63^. Moreover, miR-206, which directly inhibits PTP1B expression, can significantly reduce CM apoptosis and myocardial infarct size in rats^64^.While these data suggest a protective role for PTP1B in the heart, the precise mechanisms for how PTP1B directly impacts cardiac function have remained, in part, unclear. Indeed, most previous studies have focused only on the molecular effects of PTP1B in cardiac endothelial cells. For example, overexpression of PTP1B in bovine aortic endothelial cells dramatically inhibits VEGF-induced AKT phosphorylation^65^. Conversely, deletion of PTP1B in endothelial cells protects against cardiac hypertrophy induced by transverse aortic constriction and improves cardiac VEGF signaling and angiogenesis^45, 66^. Moreover, deletion of PTP1B in endothelial cells also promotes VEGF induced ERK activation, increasing cell proliferation and migration^46^. Here we show that CM-specific deletion of PTP1B increases both IR and VEGFR phosphorylation, validating that they are critical substrates for PTP1B in the heart ^58–60^. Consequently, CM-specific deletion of PTP1B also leads to increased downstream activity of ERK and AKT.

Activated ERK1/2 is involved in inducing a physiological hypertrophy, with enhanced contractile force and reduced fibrosis^67, 68^. However, CM-specific deletion of ERK1/2 in mice showed no reduction in pathologic or physiologic stimulus-induced cardiac growth^69^. Similarly, increased AKT activity protects against ischemia-reperfusion injury^70^. However, overt cardiac-specific overexpression of constitutively active AKT can also induce pathological cardiac hypertrophy in mice^71, 72^, suggesting that dosage of AKT is critical to the modulation of cardiac hypertrophy. Taken together, ERK and/or AKT activation in the heart may potentially have positive therapeutic effects, maintaining cardiac performance and preventing the transition to maladaptive hypertrophy and heart failure. However, this also may be dependent on a number of factors, including whether the effects are acute or chronic and whether other modulators and/or signaling pathways are involved in cardiac function. Further research is needed to fully understand the complex signaling networks involved in these processes.

AKT is also directly involved in autophagy, inhibiting this process through activation of mTOR and leading to accumulation of damaged organelles, which can contribute to the development of cardiac dysfunction. Indeed, previous studies have implicated autophagy and mitophagy in HFD-associated mitochondrial dysfunction in the heart^73, 74^. In response to short-term HFD feeding, genetic ablation of Atg7, a critical modulator of autophagy, impairs mitophagy and leads to cardiac dysfunction^74^. Here, in response to deletion of PTP1B, we observed increased AKT activity, but decreased mTOR activity and increased autophagy, which may explain, at least in part, the mechanism by which PTP1B deletion may protect against HFD-induced cardiac hypertrophy and mitochondrial dysfunction.

However, the mechanism for how PTP1B is involved in the regulation of mTOR and autophagy does not appear to be via expected canonical pathways. Our data reveals a novel PTP1B-PKM2-AMPK axis that mediates both mTOR and autophagic regulation. Our findings here are in line with recent studies in both adipose cell lines and pancreatic cancer cells, where PKM2 was found to be directly dephosphorylated by PTP1B, inducing its activity^33, 36^. We now identify a critical role for this same signaling axis in HFD-induced cardiomyopathy, showing that deletion of PTP1B decreases PKM2 activity and, thereby, protects against pathological cardiac hypertrophy. In this regard, direct inhibition of PKM2 has also demonstrated to be protective against development of right ventricular dysfunction in mice subjected to pulmonary artery banding^75^, as well as in mice with pulmonary hypertension^76^. PKM2 is also a known regulator of CM cell cycle critical for modulation of oxidative stress through anabolic signaling and activation of b-catenin^77^. It would be interesting to further investigate the direct role of PKM2 in HFD-induced cardiomyopathy and HF.

Mitochondrial dysfunction and energy imbalance are critical components of HFD-induced cardiomyopathy. One important consequence of impaired mitochondrial structure and function is increased reliance on glucose and decreased FAO^78–80^. With respect to effects of PTP1B on glycolysis, we observe that deletion of PTP1B in CMs leads to reduced steady-state levels of glycolytic intermediates and TCA intermediates, suggesting reduced activity or potentially increased metabolic flux. With regards to FAO, the directionality of the change in FAO in the heart is variable and model dependent. Thus, it might not be apparent if decreased FAO levels are causal to or a consequence of cardiac dysfunction in response to HFD. For example, reduced mitochondrial FAO is observed in patients with heart failure with preserved ejection fraction (HFpEF)^81–83^. In addition, a comprehensive multi-omics study of patients with hypertrophic cardiomyopathy (HCM) also found reduced levels of myocardial FAO intermediates, including decreased levels of acylcarnitine (AC) and reduced gene expression of carnitine palmitoyl transferase I (CPT1), a critical regulator of mitochondrial long-chain FAO^84–86^. These studies suggest that increasing FAO could be beneficial in certain contexts. Conversely, increased FAO associated with increased reactive oxygen species (ROS) in hearts from obese mice and rats, likely the consequence of a combination of increased generation of reducing equivalents by FAO, could be coupled with impaired OXPHOS function^87–89^. In a physiological context, increased ROS induced by FAO, could be self-limiting^90^, representing a short-term negative feedback mechanism as an adaptation to pathological stress imposed on the heart. In this, the negative effects of ROS might not outweigh the positive benefits of enhanced FAO. However, considerations for therapeutic modulation of FAO in hypertrophy and heart failure remain to be determined.

Though the mechanisms leading to changes in FAO are not yet fully understood, our findings suggest that increased FAO is critical for preservation of cardiac function in response to HFD; CM-specific deletion of PTP1B leads to increased FAO, enhanced levels of AC, induced mitochondrial respiration, and decreased accumulation of toxic lipids in the heart. Moreover, our data show that deletion of PTP1B in CMs restores the OXPHOS complex, increasing mitochondrial complexes I and II of the ETC, thereby increasing ATP synthesis and decreasing free radical production^91, 92^. Recent lines of evidence suggest a correlation between ROS and membrane potential (Δψ); in the case of mitochondrial disorders associated with OXPHOS dysfunction, a lower Δψ correlates with an increase in ROS production^93, 94^. Here, our results show that Δψ is increased in PTP1B^fl/fl^::ꭤMHC ^Cre/+^ CMs following HFD, suggesting decreased ROS production and preserved mitochondrial integrity.

We have also observed that CM-specific deletion of PTP1B promotes NAD^+^ synthesis through its ability to enhance AMPK-mediated metabolic signaling and NMN (the NAD^+^ precursor). NAD^+^ is a central metabolite in the salvage pathway and is involved in energy and redox homeostasis. Stimulating NAD^+^ synthesis has been shown to be beneficial against development of both diabetic cardiomyopathy^95^ and HFpEF^96^. Moreover, deletion of PTP1B leads to increased production of NAMPT, the rate-limiting enzyme in the salvage pathway. CM-specific overexpression of NAMPT has also been shown to be protective against development of HFD-induced cardiac hypertrophy and diastolic dysfunction in mice^95^. Furthermore, NAMPT overexpression restores levels of NAD^+^ and NADP^+^, protecting NDUFS4 (mitochondrial complex I subunit)-deficient mice from development of diabetic cardiomyopathy^97^. Finally, systemic NAMPT overexpression has been shown to protect mice against development of angiotensin II-induced hypertension^98^. Here, we show that increased NAD^+^ signaling in PTP1B^fl/fl^::ꭤMHC ^Cre/+^ hearts is mediated by the effects of deletion of PTP1B on PKM2 and the AMPK axis, leading to elevated mitochondrial FAO^62, 99^ and increased phosphorylation of NAMPT^100^. In line with our findings, intraperitoneal injection of NMN stimulates NAD^+^ biosynthesis in obese mice^101^, preventing development of cardiac hypertrophy^102^. Future research will be needed to determine the translational application of NAD^+^ and/or its precursors in the prevention of diabetic cardiomyopathy.

While our results herein suggest a protective cardiac effect in response to deletion of PTP1B in the heart, one recent paper suggests a different role; Coulis et al. suggest that CM-specific deletion of PTP1B in mice may be pathological, inducing a hypertrophic phenotype that is exacerbated by pressure overload^103^. Specifically, they claim that argonaute 2 (AGO2), a critical component of the RNA-induced silencing complex, is inactivated in response to CM-specific deletion of PTP1B, thereby preventing miR-208b mediated inhibition mediator complex subunit 13 (MED13) and leading to thyroid hormone-mediated pathological cardiac hypertrophy. There could be several reasons why we, and others, might have seemingly opposing results to this paper. First, inhibition of miR-208b has been previously shown to improve, not induce, cardiac dysfunction in titin-induced dilated cardiomyopathy^104^. Second, the upregulation of MED13 in the heart has a demonstrated protective function, conferring resistance to obesity^105^. Third, with respect to the effects of PTP1B directly, Coulis et al used the PTP1B^fl/fl^ mice as their control group instead of αMHC^Cre/+^; this is critical, as there is a large body of evidence indicating that expression of the αMHC-Cre transgene alone in some strains can exert a cardiac phenotype, including hypertrophy^106^. Moreover, in this regard, it is critical to use mice from the appropriate background Cre control when comparing genetic changes in mice^107, 108^. Finally, it is possible that different pathological stimuli can lead to different outcomes. For example, Byun et al. found that overexpression of NAMPT in response to pressure overload can induce heart failure through activation of Sirt1^109^, whereas, in response to HFD, Oka et al. found that NAMPT overexpression can reduce cardiac diastolic dysfunction, apoptosis and proinflammatory signaling through upregulation of NAD^+^, NADP^+^ and NADPH^95^. Exactly how PTP1B is involved in modulating various physiological vs. pathological responses in the heart remains as yet to be determined.

One other noteworthy point is that our data suggest potential sex-driven differences in the cardiometabolic alterations associated with obesity-related cardiomyopathy. It took longer for female mice to develop signs of cardiomyopathy in response to HFD feeding (20 weeks vs. 10 weeks for the males), irrespective of genetic background. In addition, the overall phenotypic observations of hypertrophy in response to HFD were less pronounced in female mice. Metabolically, however, while changes in NAD^+^ metabolites were not significantly affected in females, gene expression of the key metabolic enzymes involved in NAD^+^ synthesis were in fact elevated in both male and female PTP1B^fl/fl^::ꭤMHC ^Cre/+^ mouse hearts, suggesting that, at least at the molecular level, effects of pathological stress can be similarly attributed to both males and females. Further elucidation of potential sex-specific parallel signaling pathways involved in HFD-induced cardiomyopathy pathogenesis should, however, be considered for future studies.

Taken together, our results suggest that CM-specific deletion of PTP1B is protective and should be considered as a potential therapeutic target against development of HFD induced cardiomyopathy. Deletion of PTP1B in CMs mediates a substrate switch from glucoses to FA metabolism, protecting hearts against development of HFD-induced cardiac hypertrophy and dysfunction through mechanisms involving a novel PTP1B/PKM2/AMPK axis that is critical for the regulation of NAMPT and NAD+ biosynthesis.

## Non-standard Abbreviations and Acronyms

18S: 18S ribosomal RNA
ꭤMHC: alpha myosin heavy chain (MYH6)
ACADVL: acyl-CoA dehydrogenase very long chain
AGO2: argonaute 2
AMP: adenosine monophosphate
AMPK: AMP-activated protein kinase
ANF: atrial natriuretic factor
ATP: adenosine triphosphate
CM: cardiomyocyte
CPT1b: carnitine palmitoyl-transferase 1b
CVD: cardiovascular disease
EEF1A1: elongation factor 1-alpha 1
ERK: extracellular signal-regulated kinase
ESI: electrospray ionization
ETC: electron transport chain
FAO: fatty acid oxidation
FCCP: carbonyl cyanide-p-trifluoromethoxyphenyl-hydrazon
H&E: hematoxylin/eosin
HADHB: acetyl-CoA acyltransferase
HFD: high-fat diet
HFpEF: heart failure with preserved ejection fraction
HSL: hormone-sensitive lipase
IR: insulin receptor
IRS: insulin receptor substrate
LC3B: microtubule-associated protein 1A/1B-light chain 3B
LVIDd: end-diastolic internal dimensions of the left ventricle
LVIDs: end-systolic internal dimensions of the left ventricle
LVPW: left ventricular posterior wall thickness
MAPK: Ras–mitogen-activated protein kinase
MED13: mediator complex subunit 13
mTOR: mammalian target of rapamycin
MYH6: α-myosin heavy chain
MYH7: b-myosin heavy chain
NAD: nicotinamide adenine dinucleotide
NADPH: nicotinamide adenine dinucleotide phosphate
NAM: nicotinamide
NAMPT: nicotinamide phosphoribosyl-transferase
ND: normal diet
NDUFB8 NADH: ubiquinone oxidoreductase subunit B8
NMN: nicotinamide ribonucleotide
OCR: oxygen consumption rates
OXPHO: oxidative phosphorylation
PDH: pyruvate dehydrogenase
PI3K: phosphatidylinositol 3-kinase
PKB/AKT: protein kinase B
PKM2: pyruvate kinase muscle isozyme 2
POMC: pro-opiomelanocortin
PTP1B: protein tyrosine phosphatase
PTP1B^+/+^::ꭤMHC^Cre/+^: alpha myosin heavy chain promoter driven by expression of Cre in the background of the wildtype PTP1B allele
PTP1B^fl/fl^::ꭤMHC^Cre/+^: alpha myosin heavy chain promoter driving deletion of PTP1B specifically in cardiomyocytes
ROS: reactive oxygen species
RPL4: ribosomal protein L4
RSD: relative standard deviation
SDHB: succinate dehydrogenase subunit b
SDS-PAGE: sodium dodecyl-sulfate polyacrylamide gel electrophoresis
TCA: tricarboxylic acid
TEM: transmission electron microscope
VEGF: vascular endothelial growth factor
VEGFR: vascular endothelial growth factor receptor
WGA: wheat germ agglutinin

## Acknowledgements

We would like to thank Dr. Benjamin G. Neel (NYU Grossman School of Medicine, Laura and Isaac Perlmutter Cancer Center, New York) for his generous support in providing the PTP1B^fl/fl^ mice for these studies. Thank you to Dr. Bing Xu for his guidance on isolating primary adult cardiomyocytes. Special thanks as well to Dr. Coralie Poizat for her careful reading of our manuscript. Finally, thanks to the analytical and technical services at the State University of New York (SUNY) College of Environmental Science and Forestry for providing us the transmitted electron microscope data on our mouse hearts. We would also like to acknowledge that Figures 7 and S7 were created using BioRender.com.

## Sources of Funding

This work was supported by the National Institutes of Health (Grants R01-HL122238, R01-HL102368), the American Heart Association Transformation Grant Award (20TPA35490426), and the Masonic Medical Research Institute to M.I.K.; and by the Halfond-Weil Postdoctoral fellowship to Y.S.

## Disclosures

M.I.K. has received grant funding from Onconova Therapeutics and is a consultant for BioMarin Pharmaceutical Inc; both funding and consulting projects are independent of the work in this manuscript (no overlap).

---

**Figure S1.**
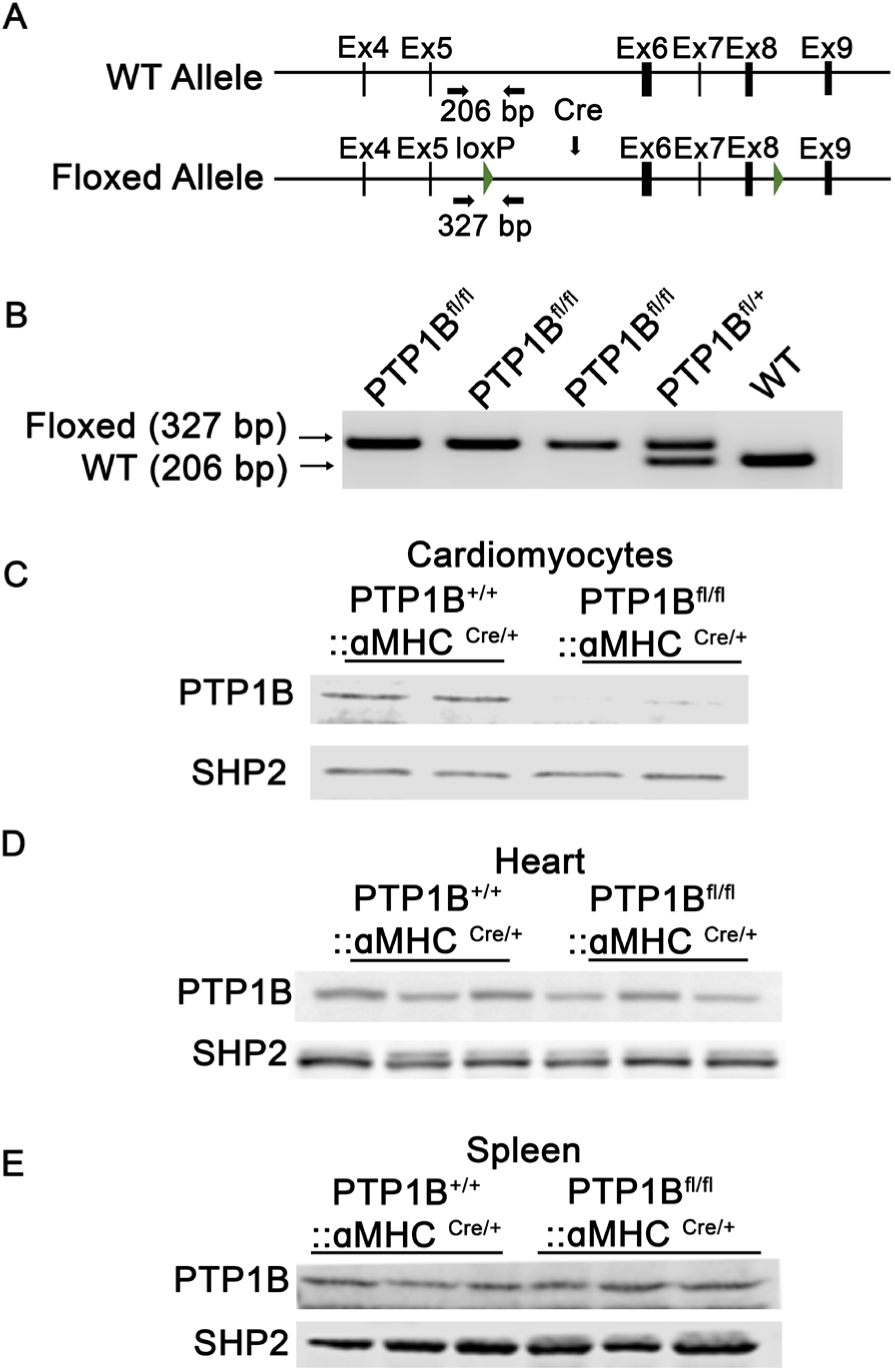
Characterization of mice with cardiac-specific deletion of PTP1B. **A.** Localization of loxP on the genome. Arrows indicate genotyping primers. **B.** Genotyping by PCR analysis. The wildtype allele has a 206-bp product while floxed allele has a product of 327 base-pair. PTP1B protein expression levels in PTP1B^+/+^::ꭤMHC^Cre/+^ and PTP1B^fl/fl^::ꭤMHC^Cre/+^ **C.** cardiomyocytes, **D.** whole heart and **E.** spleen. SHP2 expression is immunoblotted as a loading control.

**Figure S2.**
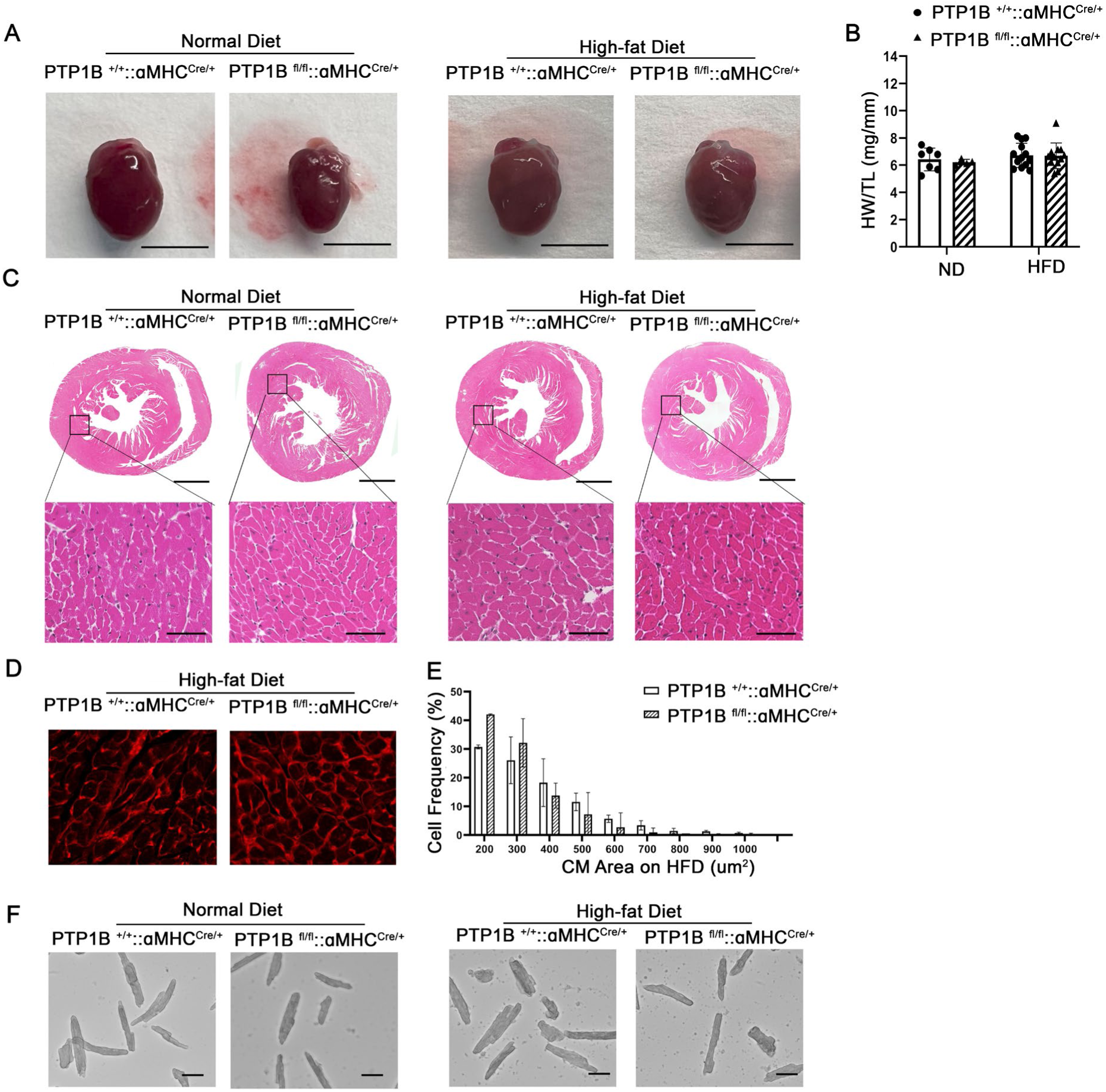
Cardiomyocyte-specific deletion of PTP1B prevents HFD-induced cardiomyopathy in female mice. **A**. Representative photographs of hearts from female PTP1B^+/+^::ꭤMHC^Cre/+^ and PTP1B^fl/fl^::ꭤMHC^Cre/+^ mice maintained on ND or HFD for 20 weeks, scale bar=50 mm. **B**. Heart weight to tibiae length ratios from female control or PTP1B^fl/fl^::ꭤMHC^Cre/+^ mice on ND or HFD. n= 8-12/group. * p<0.05 and **p<0.01 from two-way ANOVA. **C**. Representative H&E whole heart cross-sections from PTP1B^+/+^::ꭤMHC^Cre/+^ and PTP1B^fl/fl^::ꭤMHC ^Cre/+^ female mice maintained on ND or HFD for 20 weeks. Upper panel scale bar=1 mm; Lower panel, scale bar=50 µm. **D**. Wheat germ agglutinin (WGA) staining (red) from heart cross-sections from control and PTP1B^fl/fl^::ꭤMHC^Cre/+^ female mice fed a ND or HFD for 20 weeks. Scale bar= 20 µm. **E**. Frequency distribution of CM area from control and PTP1B^fl/fl^::ꭤMHC^Cre/+^ female mice fed a ND or HFD. N=3 mice per group, with at least 1x10^3^ cell counts per heart. Data are represented as mean ± SEM. * p<0.05 based on Student’s *t*-test. **F**. Representative photomicrograph of ventricular myocytes isolated from PTP1B^+/+^::ꭤMHC^Cre/+^ and PTP1B^fl/fl^::ꭤMHC^Cre/+^ female mice maintained on ND or on HFD. Scale bar= 50 µm.

**Figure S3.**
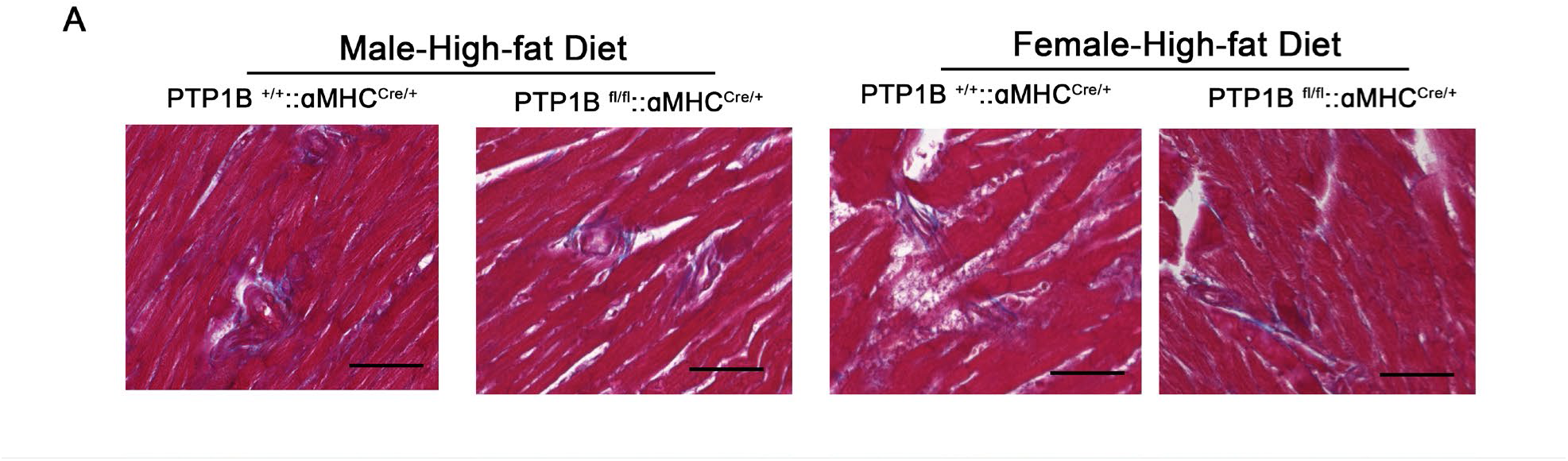
Effects of deletion of PTP1B on fibrosis. **A**. Representative Masson’s trichrome-stained section of hearts from male and female PTP1B^+/+^::ꭤMHC^Cre/+^ and PTP1B^fl/fl^::ꭤMHC^Cre/+^ mice maintained on HFD. Scale bar= 50 µm.

**Figure S4.**
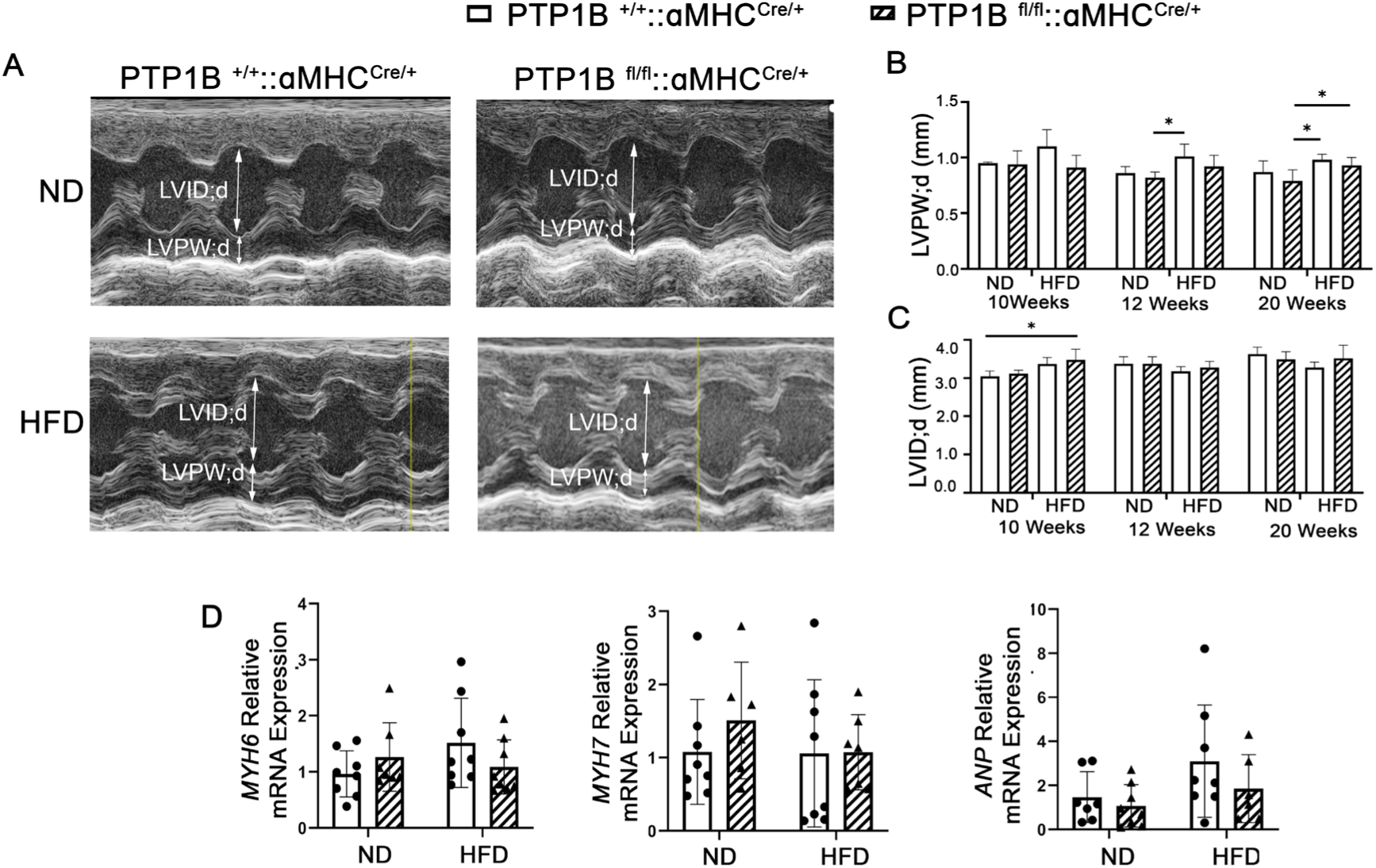
Cardiomyocyte-specific PTP1B deletion preserves cardiac function in response to HFD in female mice. **A**. Representative echocardiography of PTP1B^+/+^::ꭤMHC^Cre/+^ and PTP1B^fl/fl^::ꭤMHC^Cre/+^ female mice fed a ND or a HFD for up to 20 weeks. A two-headed arrow indicates left ventricular chamber size or left ventricular wall thickness. LVID;d= left ventricular internal dimension in diastole. LVPW;d=left ventricular posterior wall thickness in diastole. Quantification of **B.** LVPWd and **C.** LVIDd (n=10-15/group). **D.** Real-time q-PCR analysis of hypertrophy-related gene expression (*MYH6, MYH7, and ANP*) in CM mRNA isolated from either PTP1B^+/+^::ꭤMHC^Cre/+^ or PTP1B^fl/fl^::ꭤMHC^Cre/+^ female mice maintained on ND or HFD for 20 weeks. The ratio of ΔΔCT was analyzed using *18S* and eukaryotic elongation factor-1 (*Eef1*) mRNA as housekeeping controls, and HFD results were compared to ND from the same genotype. Samples were obtained from 6-7 mice per group and each sample was assessed in technical triplicate. Data in graphs represent mean ± SEM; *p<0.05, ** p<0.01, ***p<0.001, and all p values were derived from ANOVA with Bonferroni post-test when ANOVA was significant.

**Figure S5.**
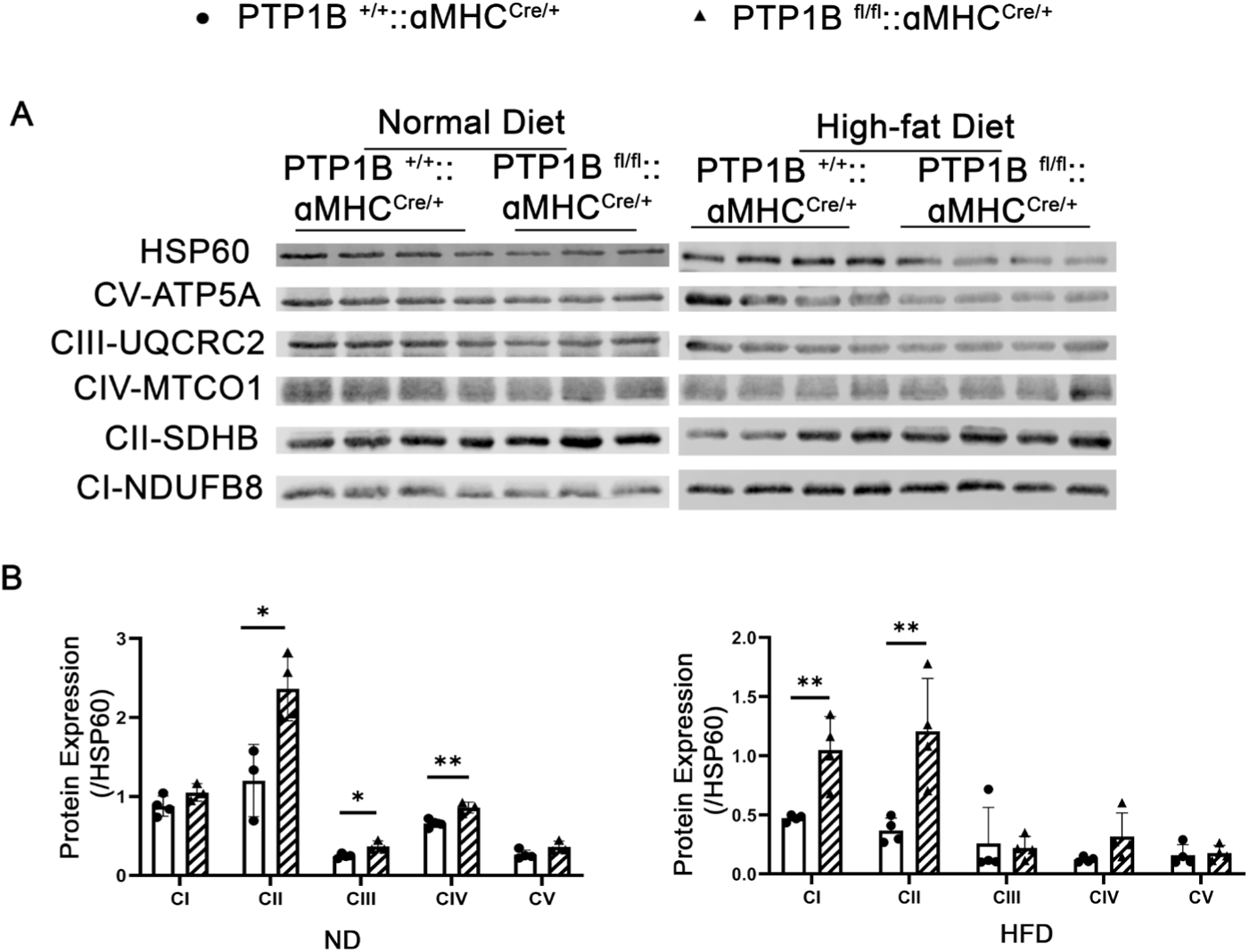
Cardiomyocyte-specific deletion of PTP1B increases electron transport chain subunit levels in HFD female mice. **A**.Western blot and the **B.** quantitative assessment of OXPHOS mitochondrial complexes garnered from female PTP1B^+/+^::ꭤMHC^Cre/+^ and PTP1B^fl/fl^::ꭤMHC^Cre/+^ hearts on HFD for 20 weeks (n=6 mice per group). Immunoblot utilized an antibody cocktail that recognizes the five mitochondrial oxidative phosphorylation complexes. *p<0.05, **p<0.01 using one-way ANOVA.

**Figure S6.**
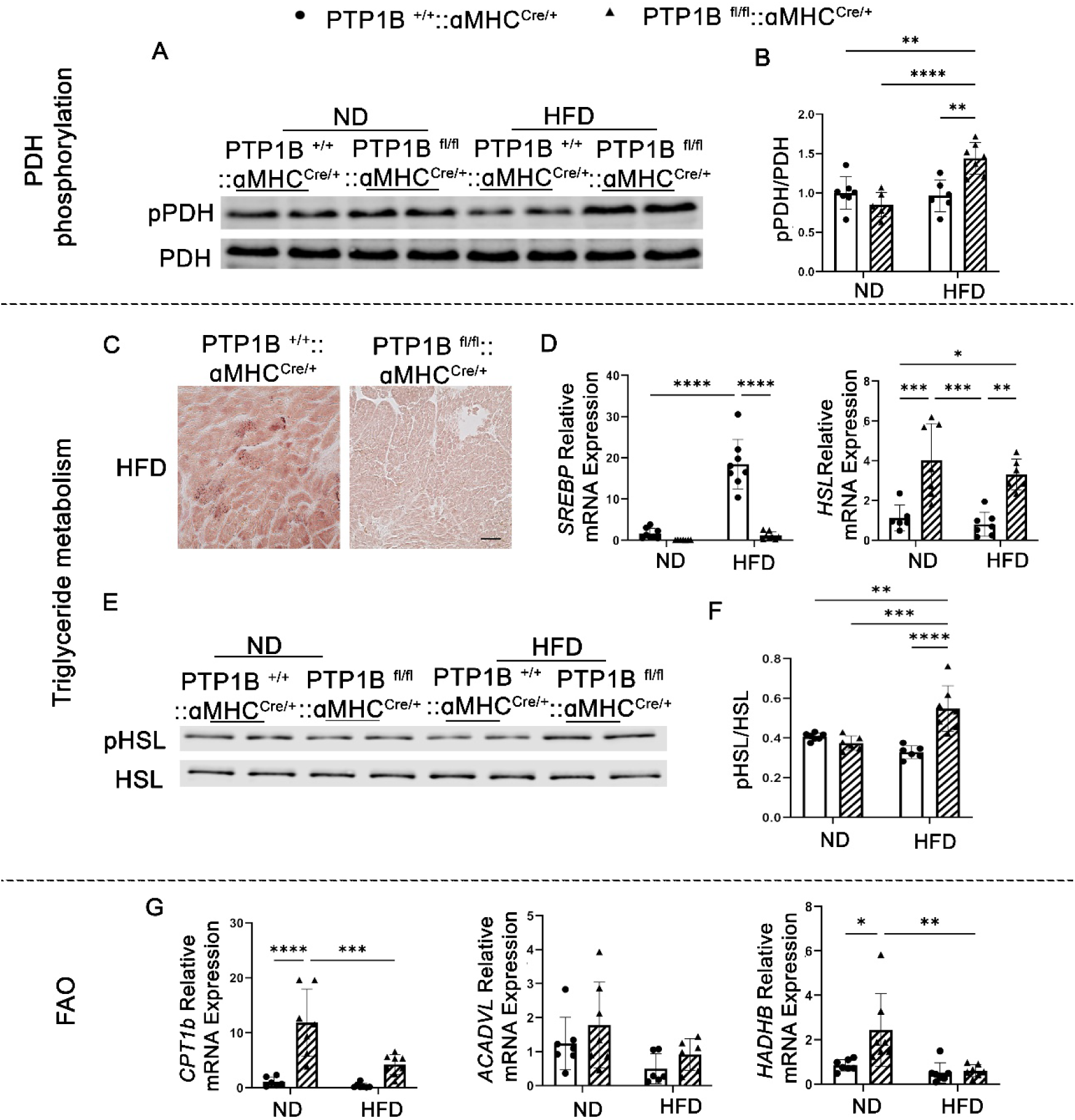
Triglyceride mobilization and fatty acid oxidation are increased in female PTP1B^fl/fl^::ꭤMHC^Cre/+^ hearts. **A.** Representative western blot of heart lysates immunoblotted with phospho-PDH (Ser 293) or PDH antibodies. **B.** Quantification of pPDH/total PDH ratios. N=6-7 mice/group. **C.** Oil Red O staining lipid droplets in frozen female heart sections from both PTP1B^+/+^::ꭤMHC^Cre/+^ and PTP1B^fl/fl^::ꭤMHC^Cre/+^ mice fed with HFD for 20 weeks. Scale bar= 50 µm. **D**. Gene mRNA expression analysis (real-time PCR) of lipogenic transcription factors *SREBP* and *HSL* in CMs isolated from female PTP1B^+/+^::ꭤMHC^Cre/+^ and PTP1B^fl/fl^::ꭤMHC^Cre/+^. The ratio of ΔΔCT was analyzed using *18S* and eukaryotic elongation factor-1 (*Eef1*) mRNA as housekeeping controls, and HFD results were compared to ND from the same genotype. Samples were obtained from 7-9 mice per group and each sample was assessed in technical triplicate. Data in graphs represent mean ± SEM; *p<0.05, ** p<0.01, ***p<0.001, and all p values were derived from ANOVA with Bonferroni post-test when ANOVA was significant. **E**. Representative western blotting analysis and **F.** quantitative analysis of HSL protein expression in heart lysates isolated from female PTP1B^+/+^::ꭤMHC^Cre/+^ or PTP1B^fl/fl^::ꭤMHC^Cre/+^ mice fed HFD for 20 weeks. N=6-7 mice/group. **G**. Gene expression of fatty acid oxidation genes *CPT1b*, *ACADVL*, and *HADHB* in CMs isolated from female PTP1B^+/+^::ꭤMHC ^Cre/+^ or PTP1B^fl/fl^::ꭤMHC ^Cre/+^ mice fed with either ND or HFD for 20 weeks. The ratio of ΔΔCT was analyzed using *18S* and eukaryotic elongation factor-1 (*Eef1*) mRNA as housekeeping controls, and HFD results were compared to ND from the same genotype. Samples were obtained from 7-9 mice per group and each sample was assessed in technical triplicate. Data in graphs represent mean ± SEM; *p<0.05, ** p<0.01, ***p<0.001, and all p values were derived from ANOVA with Bonferroni post-test when ANOVA was significant. Statistical significance was determined with 2-way ANOVA. *p<0.05,**p<0.01,***p<0.001.

**Figure S7.**
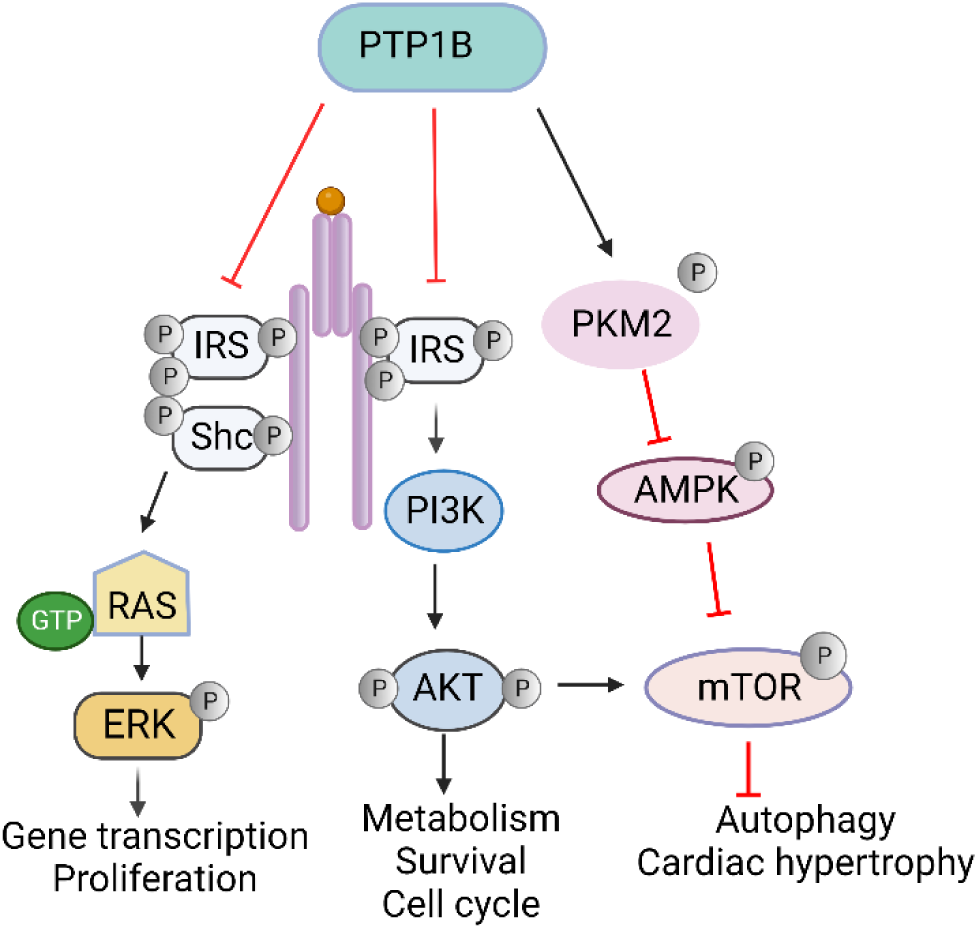
PTP1B signaling in autophagy and cardiac hypertrophy. PTP1B acts as a negative regulator of the insulin signaling pathway. PTP1B dephosphorylates insulin receptor and/or insulin receptor substrate (IRS), inactivating downstream pathways that include PI3K/AKT/mTOR and ERK-MAPK signaling. PTP1B also dephosphorylates PKM2, leading to its activation, thereby inhibiting downstream signaling to AMPK and mTOR.

**Figure S8.**
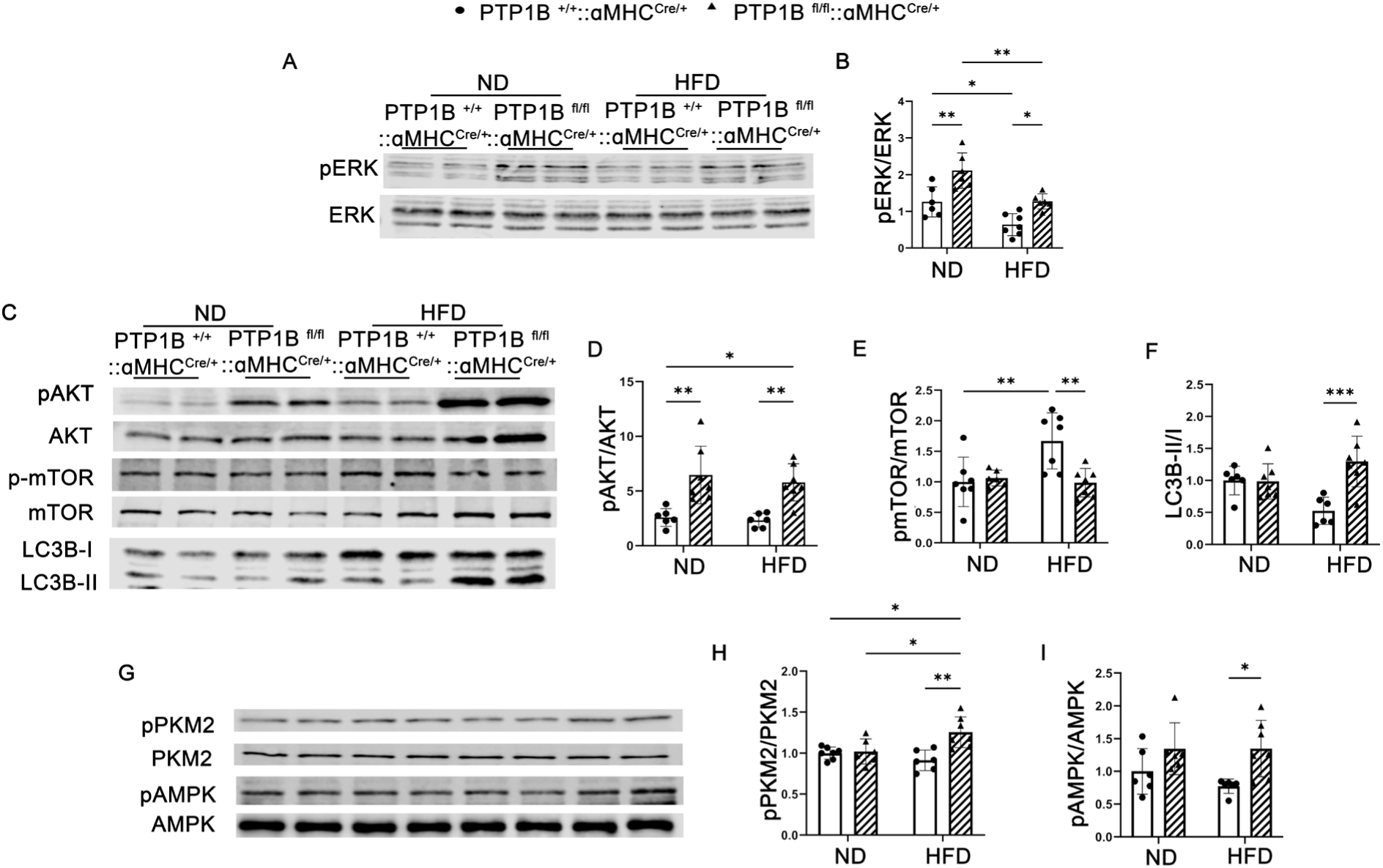
Loss of PTP1B in CMs induces activation of ERK, AKT, PKM2 and AMPK. **A**. Representative western blot of phospho and total ERK in cardiac extracts isolated from female PTP1B^+/+^::ꭤMHC ^Cre/+^ or PTP1B^fl/fl^::ꭤMHC ^Cre/+^ mice fed a ND or HFD. **B.** Quantificatative analyses of the ratio of ERK phosphorylation to total ERK expression generated from three independent experiments, n=6-8 hearts/group. **C-F.** Representative western blots and quantitative analysis of total and phospho-AKT (Ser 473), total and phospho-mTOR (Ser 248), as well as the autophagy markers, LC3I and LC3II, in cardiac lysates isolated from female PTP1B^+/+^::ꭤMHC^Cre/+^ or PTP1B^fl/fl^::ꭤMHC^Cre/+^ mice fed ND or HFD. N=6-8 hearts/group. **G-I**. Representative western blots and quantitative analysis of heart lysates immunoblotted with phospho-PKM2 (Tyr105), as well as total and phospho-AMPK (Thr 172). n=6-8 hearts/group.

**Figure S9.**
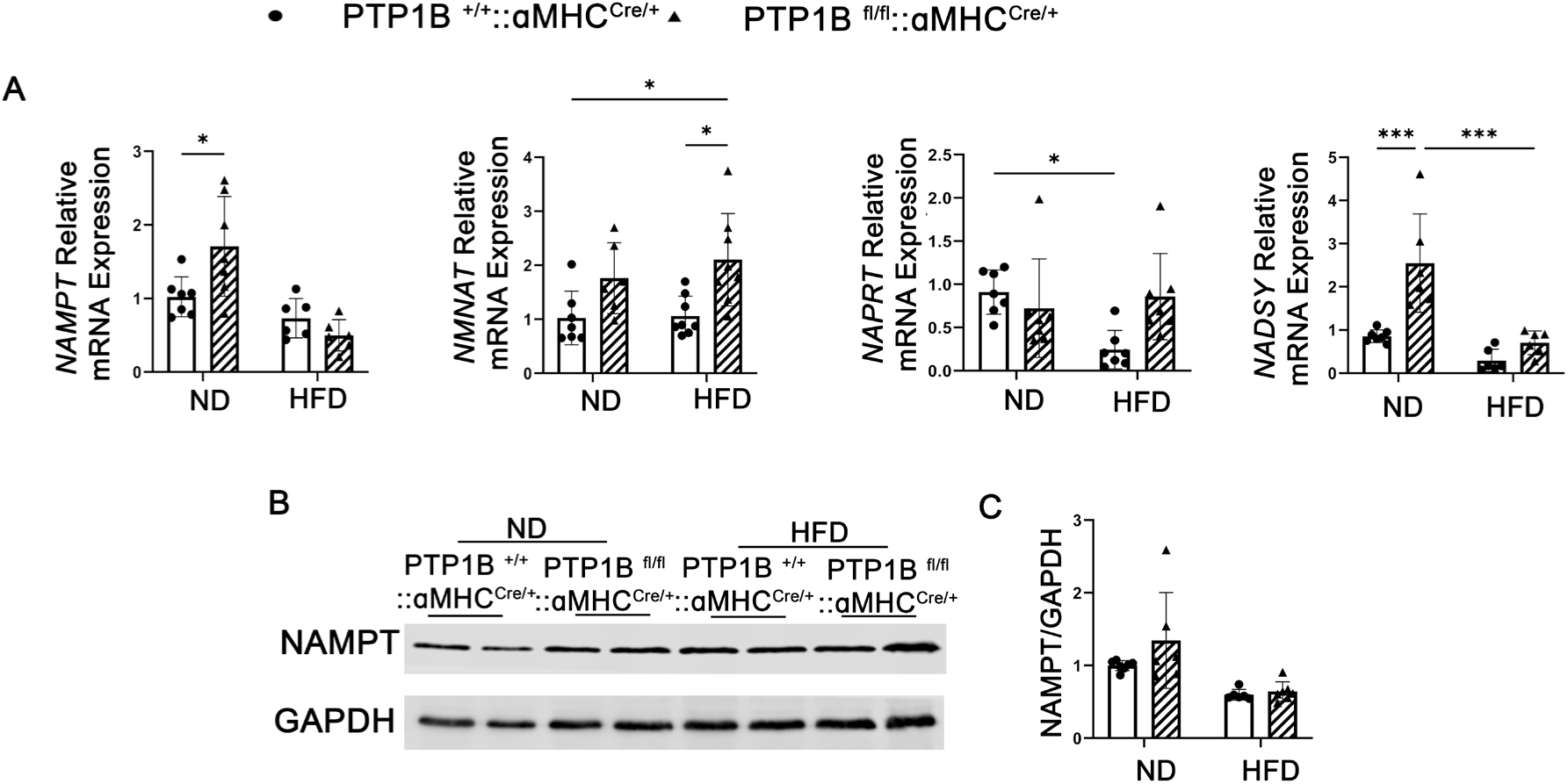
Genes involved in NAD^+^ synthesis are upregulated in female PTP1B^fl/fl^::ꭤMHC^Cre/+^ mice. **A.** NAD^+^ salvage synthesis pathway critical genes *NAMPT*, *NMNAT*, *NAPRT*, *NADSYN* were measured by quantitative real-time PCR in CMs from female PTP1B^+/+^::ꭤMHC^Cre/+^ and PTP1B^fl/fl^::ꭤMHC^Cre/+^ mice fed ND or HFD for 20-weeks . The ratio of ΔΔCT was analyzed using *18S* and eukaryotic elongation factor-1 (*Eef1*) mRNA as housekeeping controls, and HFD results were compared to ND from the same genotype. Samples were obtained from 7-9 mice per group and each sample was assessed in technical triplicate. Data in graphs represent mean ± SEM; *p<0.05, ** p<0.01, ***p<0.001, and all p values were derived from ANOVA with Bonferroni post-test when ANOVA was significant. Statistical significance was determined with 2-way ANOVA. *p<0.05, ** p<0.01, ***p<0.001. **B-C**. Representative western blots and quantitative analyis of heart lysates immunoblotted with NAMPT and GAPDH (control for loading). N=6-8 hearts/group).Statistical significance was determined with 2-way ANOVA. *p<0.05, ** p<0.01,***p<0.001.

**Table S1.**
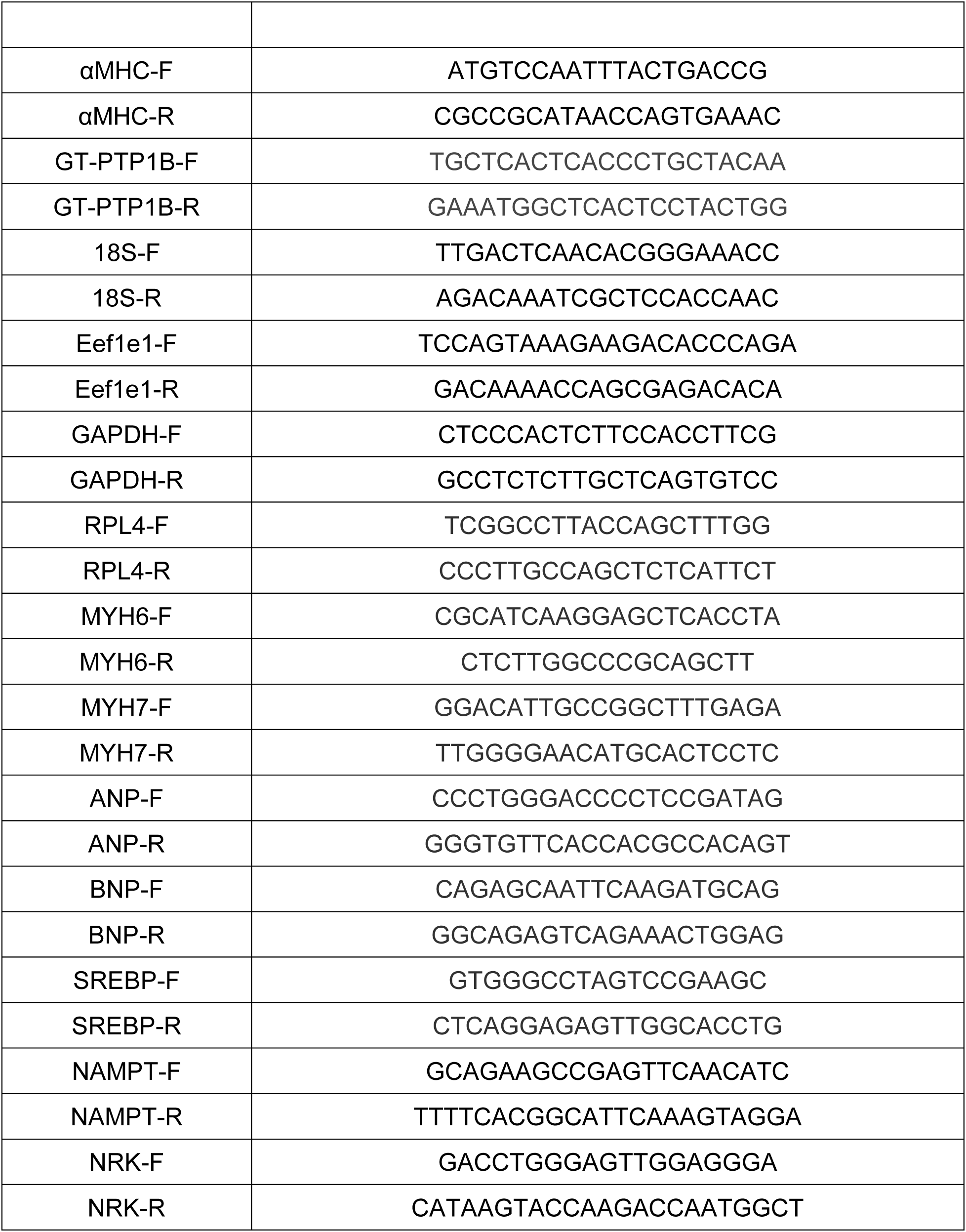
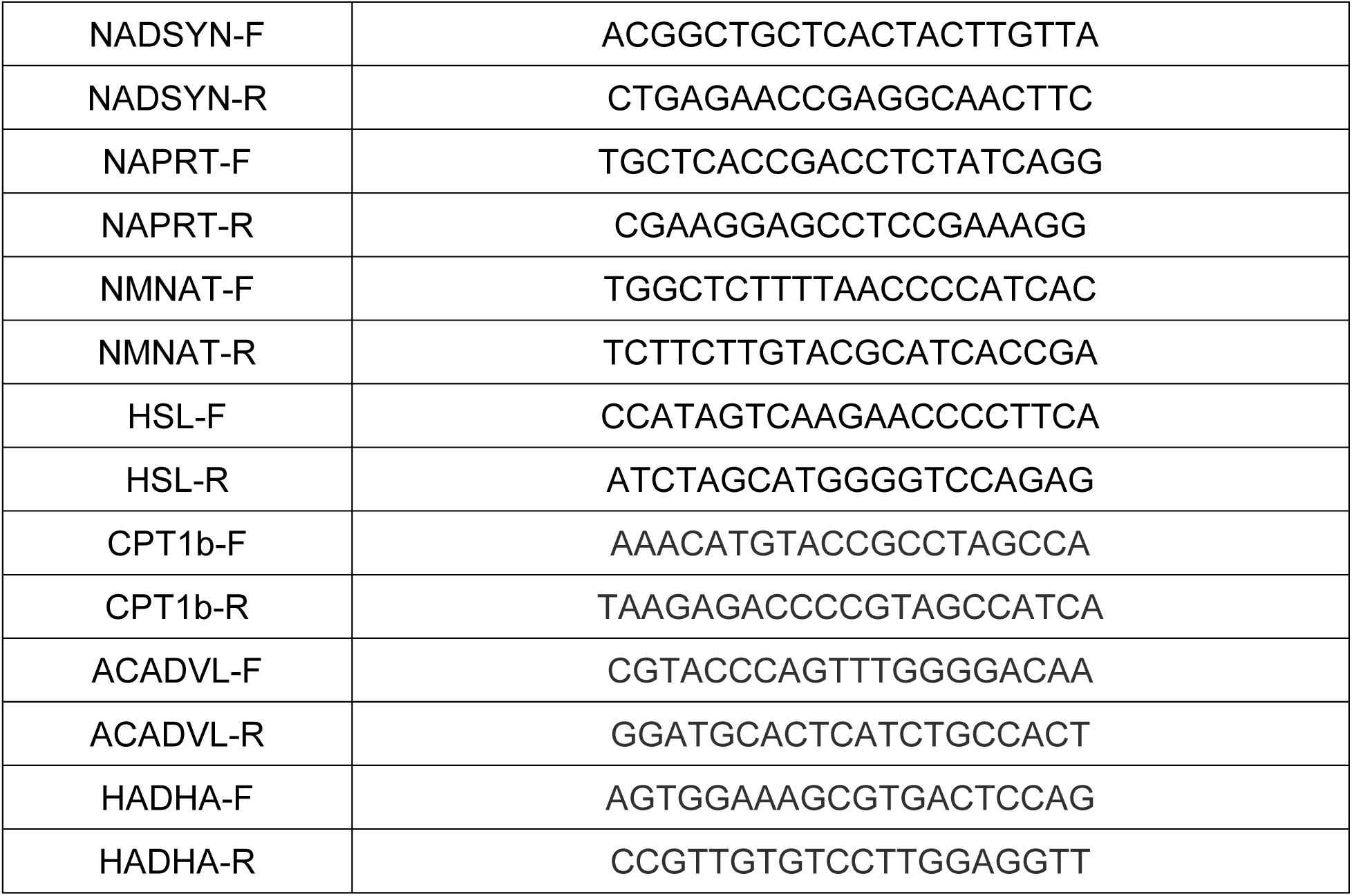
Primer sequence.

**Table S2.** Mendelian ratios of newborn mice from crossing of PTP1B^fl/fl^::ꭤMHC ^Cre/+^ and PTP1B^fl/fl^.

| PTP1B<sup>fl/fl</sup>::αMHC<sup>Cre/+</sup> X PTP1B<sup>fl/fl</sup> |  |  |  |
| --- | --- | --- | --- |
|  | Observed number/ratio | Expected number/ratio | P-value |
| PTP1B<sup>fl/fl</sup> | 469/52.6% | 446/50% | n.s. |
| PTP1B<sup>fl/fl</sup>::αMHC<sup>Cre/+</sup> | 423/47.4% | 446/50% | n.s. |

- Mice were genotyped for at 3-weeks of age after weaning - P-values of the mendelian ratios are calculated with the Chi-square test and are either significant if p<0.05 or not significant(n.s.)

**Table S3.** Measurements of isolated cardiomyocytes from female mice.

|  | ND |  | HFD |  |
| --- | --- | --- | --- | --- |
|  | PTP1B<sup>+/+</sup> ::<br>αMHC<sup>Cre/+</sup> | PTP1B<sup>fl/fl</sup> ::<br>αMHC<sup>Cre/+</sup> | PTP1B<sup>+/+</sup> ::<br>αMHC<sup>Cre/+</sup> | PTP1B<sup>fl/fl</sup> ::<br>αMHC<sup>Cre/+</sup> |
| Length<br>(μm) | 112.6±11.6 | 119.4±13.1 | 128.4±8.5 | 122.4±10.9 |
| Width<br>(μm) | 24.0±3.5 | 24.9±5.9 | **36.8±4.8\*\*** | 26.6±3.6 |
| Avg Area<br>(μm<sup>2</sup>) | 2384.1±88.5 | 2552.3±147.2 | **3266.9±145.6\*\*** | 2672.9±191.2 |

**Table S4.**
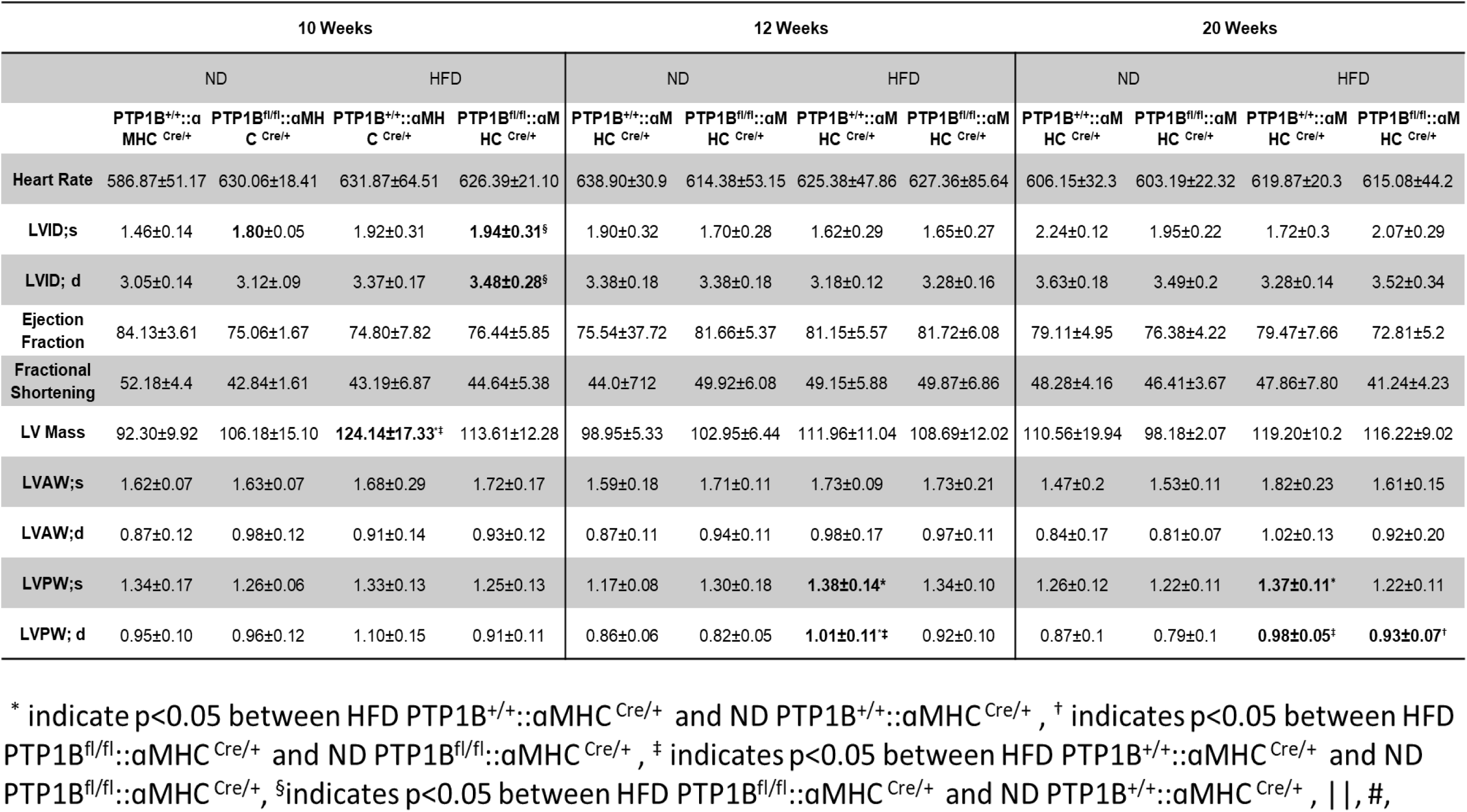
Echocardiographic analyses of female mice treated with ND or HFD for 8-12 weeks.

**Table S5.** Statistical summary of heart tissue metabolite profile differences between control and PTP1B^fl/fl^::ꭤMHC^Cre/+^ mice fed HFD.

| Statistical comparison |  |  |
| --- | --- | --- |
|  | Male | Female |
| **ANOVA** | <u>PTP1B<sup>fl/fl</sup>::αMHC<sup>Cre/+</sup> HFD</u> | <u>PTP1B<sup>fl/fl</sup>::αMHC<sup>Cre/+</sup> HFD</u> |
| **contrasts** | PTP1B<sup>+/+</sup>::αMHC<sup>Cre/+</sup> HFD | PTP1B<sup>+/+</sup>::αMHC<sup>Cre/+</sup> HFD |
| **Total biochemicals**<br>**P ≤ 0.05** | 77 | 64 |
| **Biochemicals** | ↑18\|59↓ | ↑29\|35↓ |

## Notes

### Competing Interest Statement

The authors have declared no competing interest.

